# Genomic and functional adaptations in guanylate-binding protein 5 (GBP5) highlight specificities of bat antiviral innate immunity

**DOI:** 10.1101/2025.02.11.637683

**Authors:** Amandine Le Corf, Sarah Maesen, Clara Loyer, Juan Manuel Vazquez, M. Elise Lauterbur, Lucas Sareoua, Genevieve Gray-Sandoval, Andrea Cimarelli, Carine Rey, Peter H Sudmant, David Enard, Lucie Etienne

**Author notes:** Equal contribution (Juan Manuel Vazquez, M. Elise Lauterbur). Department of Biology, University of Vermont, Burlington, VT, USA.

## Abstract

Bats are asymptomatic reservoirs of several zoonotic viruses. This may result from long-term coevolution between viruses and bats, that have led to host adaptations contributing to an effective balance between strong antiviral responses with innate immune tolerance. To better understand these virus-host interactions, we combined comparative transcriptomics, phylogenomics and functional assays to characterize the evolution of bat innate immune antiviral factors. First, we stimulated the type I interferon immune pathway in *Myotis yumanensis* primary cells and identified guanylate-binding protein 5 (GBP5) as the most differentially expressed interferon-stimulated gene (ISG). Phylogenomic analyses showed that bat GBP5 has been under strong episodic positive selection, with numerous rapidly evolving sites and species-specific gene duplications, suggesting past evolutionary arms races. Functional tests on GBP5 orthologs from ten bat species covering the >60 million years of Chiroptera evolution revealed species- and virus-specific restrictions against RNA viruses (retrovirus HIV, and rhabdoviruses European bat lyssavirus and VSV), which are typical signatures of adaptations to past viral epidemics. Interestingly, we also observed a lineage-specific loss of the GBP5 prenylation motif in the common ancestor of *Pipistrellus* and *Eptesicus* bats, associated with different GBP5 subcellular localization and loss of antiviral functions. Resurrection of the ancestral prenylation motif in *Eptesicus fuscus* GBP5 rescued its subcellular localization, but not the complete antiviral activities, suggesting that additional determinants are necessary for the antiviral restriction. Altogether, our results highlight adaptations that contribute to bat specific immunity and provide insights into the functional evolution of antiviral effector GBP5.

**Key Findings:**

- GBP5 is the most differentially expressed gene upon type I interferon stimulation of *Myotis yumanensis* primary cells.
- Bat GBP5 has evolved under strong episodic genomic and genetic diversification, including early stop codon leading to protein truncation and loss of prenylation motif.
- GBP5 diversification in bats impacts their subcellular localization and antiviral functions.
- Bat GBP5s exhibit species- and virus-specificity in their ability to inhibit infectivity of viral particles, bearing glycoproteins from retroviral HIV, vesicular stomatitis virus and European bat lyssavirus-1.
- Resurrection of the ancestral prenylation motif in a bat GBP5 rescues its subcellular localization but not its full antiviral activity

## Introduction

The mammalian antiviral innate immune system has been shaped by ancient conflicts with pathogenic viruses, which must constantly adapt to evade or antagonize host antiviral responses^1,2^. These past viral epidemics exerted selective pressures favoring genomic adaptations that naturally arose in the host and were a benefit against pathogenic viruses, either capable of inhibiting virus replication or limiting the development of pathogenesis. Over evolutionary time, host-virus arms races have left signatures of adaptations in host genomes, such as site-specific positive selection by non-synonymous mutations, gene duplications or indels^1^. Past host adaptations may provide modern immune specificities against existing viruses. These immune characteristics may be important in the context of cross-species transmissions and in shaping reservoir host species, which asymptomatically host viruses.

Among mammals, bats are reservoirs of a wide range of viruses, some of which are pathogenic to other species and have been the source of zoonotic outbreaks in humans^3^. With over 1,400 species spread across all continents and over 60 million years of divergence, bats have a long and diverse evolutionary history with viral infections. This long co-evolution has contributed to shaping their immune system and led to specific adaptations, which may participate in their efficient control of viral infections. Bats have altered expression and function of inflammasome components and other innate immune sensors, including the cellular receptors NLRP3^4^, PYHIN^5^ and AIM2. For example, the loss of inflammasome activation in some bat species may reduce the deleterious excess of inflammatory responses upon viral infections and may be protective. Despite this enhanced tolerance, their antiviral defenses appear strong and diversified. Several bat antiviral effectors, particularly from the interferon (IFN) innate immune pathway, present signatures of strong species-specific positive selection and gene duplications, which may be reminiscent of past host-pathogen arms races. This is for example the case for the Guanylate-binding protein (GBP) family, which encodes effectors against bacteria, parasite and viral infections^6^, and has undergone gene duplications/losses and site-specific positive selection during Chiroptera evolution^7^. However, no functional study has characterized the potential drivers or consequences of GBP diversifications yet. In fact, amongst the thousands of mammalian ISGs, only few studies combined evolutionary and functional approaches in bats to formally reveal their adaptations to past viral epidemics. Studies on protein kinase R (PKR)^8^, the Mx family of guanosine triphosphatases (e.g. Mx1)^9^, interferon-induced transmembrane protein 3 (IFITM3)^10,11^, or ISG15^12^ showed evidence of adaptive duplications, splice variants or site-specific adaptations in response to DNA and RNA viral infections. Yet, because of the lack of genomic and cellular tools, evolutionary and functional understanding on the antiviral innate immune response in bats remains poorly investigated.

We previously identified strong signals of positive selection in antiviral immune factors of the *Myotis* bat lineage^13^. Here, we therefore derived primary cells from three *Myotis yumanensis* individuals and performed a transcriptomic analysis of their IFN innate immune response. This allowed us to identify guanylate-binding protein 5 (GBP5) as the most differentially up-regulated interferon-stimulated gene (ISG) in *Myotis yumanensis* primary cells.

Human GBP5 is an IFN-inducible antiviral factor that restricts a broad spectrum of intracellular pathogens^14,15^, including viruses such as HIV-1 (Human immunodeficiency virus 1), SARS-CoV-2 (severe acute respiratory syndrome coronavirus 2), or bat lyssavirus EBLV-1 (European Bat LyssaVirus 1)^16–18^. GBP5 notably targets viral glycoprotein processing, leading to the release of new virions with immature glycoproteins, thereby reducing virions’ infectivity^19^. Although the exact mechanisms are still unknown and debated^19–21^, this viral restriction appears dependent on human GBP5 localization at the trans-Golgi network (TGN), thanks to its prenylation at a CaaX motif in the C-terminal end of the protein^18^. We further reasoned that studying natural variations in this antiviral effector during mammalian evolution would help understand its functions.

We therefore combined phylogenomic and functional approaches to determine bat GBP5 adaptation and function against modern viruses. We found that bat GBP5 has been under strong episodic positive selection and experienced gene duplication, suggesting important past evolutionary arms races. Functional tests on GBP5 from a large panel of ten bat species revealed species-specificity of restriction against viral particles bearing glycoproteins of diverse RNA viruses: HIV-1, VSV (vesicular stomatitis virus) and EBLV-1. Interestingly, we also found lineage-specific loss of the prenylation motif, correlating with intracellular and antiviral phenotypic variations. Resurrection of the ancestral GBP5 prenylation motif rescued the subcellular localization, but was not sufficient for complete antiviral activity. These results highlight adaptations, possibly driven by past viral epidemics, that contribute to bat immunity and result in specific modern host-virus restrictions.

## Results

### Transcriptomics of primary cells from three *Myotis yumanensis* individuals after immune stimulation unveil GBP5 as a top interferon stimulated gene

To better characterize cell-autonomous innate immunity in bats, we performed a transcriptomic approach in primary cells from three individuals of *Myotis yumanensis* under type I interferon (IFN) stimulation **(Fig. 1A-C)**. Briefly, primary fibroblast-like cells were derived from wing punches of three *M. yumanensis* bats in the US Southwest (see Methods). Following previous studies on the IFN stimulation of bat cell lines^22–24^, we stimulated the primary cells with, or without, universal type I IFN for 6 hours. We then isolated total cellular RNA for sequencing (RNAseq) and subsequent bioinformatic analyses. We used Salmon^25^ to quantify *M. yumanensis* gene expression. Through pairwise comparisons with DESeq2^26^, we identified genes that were significantly upregulated (n=1036) and downregulated (n=859) by a log2 fold change (FC) ≥ 2 and with an adjusted p value < 0.05, and which we defined as significantly responding to type I-IFN **(Fig. 1D).** Amongst the significantly up-regulated genes, we found several innate immune factors, indicating that I-IFN treatment led to an efficient activation of the innate immune responses in these primary cells. Specifically, among the most differentially expressed genes, we identified some involved in the innate immune signaling pathways (e.g. IRF7, IFIH7), cytokines (e.g. CXCL10) and antiviral ISGs, such as ISG15, ISG20, OAS1, OASL, Mx2 or RSAD2/Viperin **(Fig. 1D, S1).** Notably, GBP5 was the most differentially expressed ISG in *M. yumanensis* primary cells compared to the unstimulated control, an interesting finding in light of the broad antiviral activities recently ascribed to this factor.

**Figure 1.**
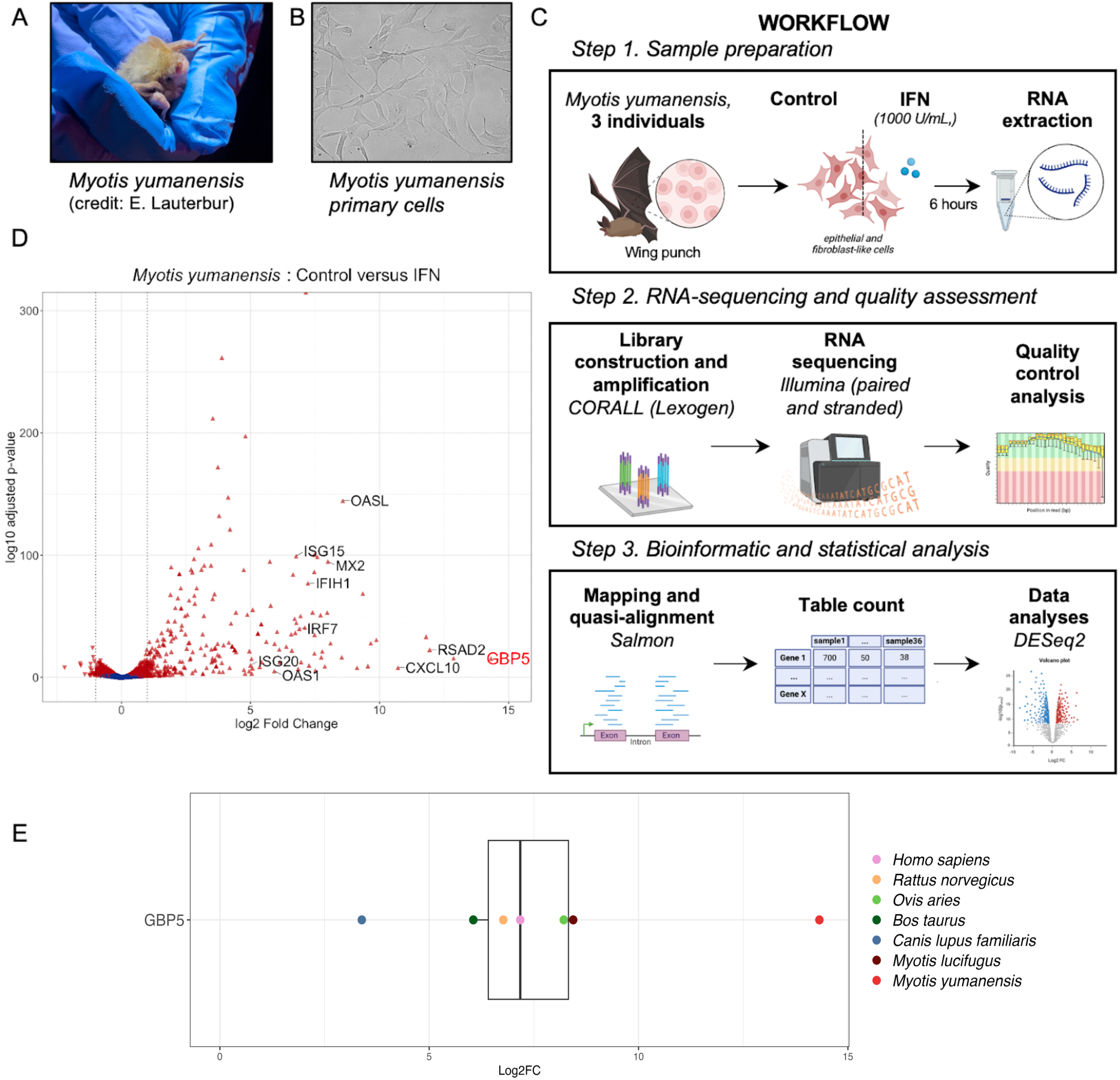
GBP5 is the most upregulated ISG in *Myotis yumanensis* primary cells upon interferon immune stimulation. A, Photo of a *Myotis yumanensis* bat at a sampling site (Credit: Elise Lauterbur). B, Primary cell line derived from biopsy wing punches. C, Schematic of the workflow from bat sampling to RNA sequencing analyses. Bat primary cell lines were similarly derived from three *M. yumanensis* individuals and treated with, or without, universal type I IFN to trigger ISG expression. Total RNA was extracted and sequenced, followed by analyses to identify differentially expressed genes between stimulated cells compared to control cells. D, Volcano plot representing the differential gene expression between untreated and IFN-stimulated cells. Genes significantly (p<0.05) differentially expressed (DE) are colored in red. Several known ISGs are highlighted on the graphic. GBP5 is the most upregulated ISG. E, Comparative analysis of differential gene expression (Log2FC, Log 2 Fold Change) upon type I IFN treatment of GBP5 in *M. yumanensis* cells with six mammalian species cells in similar experimental setup (^22^; details in Methods).

To decipher whether this was typical of mammalian GBP5 expression upon interferon stimulation, we performed a comparative analysis. We used publicly available data from a transcriptomic study defining the interferome of cells among mammalian species treated in similar conditions^22^. We found that GBP5 was also an ISG in six other mammalian species **(Fig. 1E).** Yet, the upregulation of GBP5 transcripts in *Myotis yumanensis* primary cells was remarkably high compared to the others, including its closest related species with transcriptomic information, *Myotis lucifugus* (1.5-fold higher; **Fig. 1E**).

### GBP5 has evolved under episodic positive selection during mammalian evolution

Antiviral factors that are important *in vivo* have often evolved under recurrent positive selection, as a consequence of their engagement in molecular arms-races with pathogenic viruses^2,27^. We therefore characterized the evolutionary history of GBP5 in mammals, using phylogenetic and positive selection analyses. First, we retrieved GBP5 homologous sequences in mammalian genomes from public databases using human GBP5 as a query (details in Methods). We further obtained *de novo* sequences of bat GBP5s from closely related *Myotis* species that we sampled in the wild and from a cell line of *Eptesicus fuscus* (see Methods). In total, we analyzed 348 nucleotide sequences of mammalian orthologous GBP5s from the main orders: Chiroptera (n=55), Rodentia (n=80), Primates (n=37), Perissodactyla (n=9), Artiodactyla (n=97), Lagomorpha (n=10) and Carnivora (n=51), as well as other mammalian sequences that were used as outgroups: Proboscidea (n=3), Pilosa (n=1), Cingulata (n= 4) and Xenarthra (n=1). We then aligned the GBP5 coding sequences with PRANK (codon alignment,^28^) and performed phylogenetic analyses with IQ-TREE^29^. The GBP5 gene tree generally followed the mammalian species tree **(Fig. 2A),** with some exceptions due to gene duplications/losses. For example, we confirmed an ancient duplication event in Lagomorpha^30^ and the loss of GBP5 in primate Old World Monkeys^31,32^. We also observed an important diversification for rodents and bats with duplication events, indels and high variability in sequences **(Fig. 2A, available alignments).**

**Figure 2.**
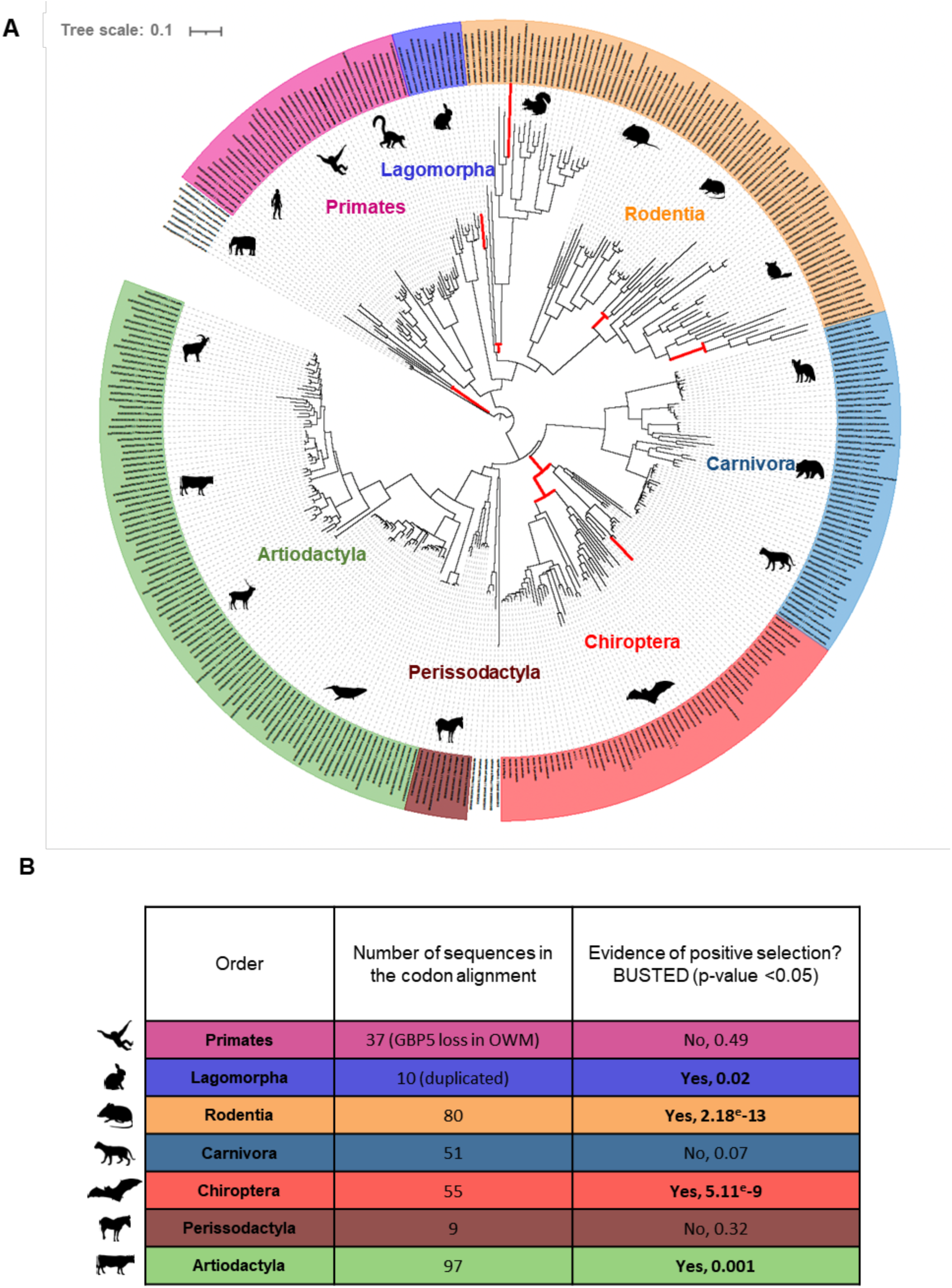
Mammalian GBP5 has evolved under episodic positive selection, with strong diversification in rodents and bats. A, Phylogenetic analysis of GBP5 across mammals. The phylogeny was built from a PRANK^28^ codon alignment of 348 GBP5 homologous sequences. Maximum likelihood phylogenetic tree was built with IQ-TREE^29^ with 1000-bootstrap replicates for statistical support (see Data availability). Branches under significant positive selection (p-value <0.05) assigned by aBSREL are thickened and in red. The scale bar indicates the number of substitutions per site. B, Evidence of positive selection in GBP5 from several mammalian orders. Positive selection analyses were performed with BUSTED^34^ using a PRANK codon alignment for each mammalian order. OWM, Old world monkeys. Species silhouettes are from https://www.phylopic.org.

To identify whether and when episodic positive selection of GBP5 has occurred during mammalian evolution, we used a branch-specific method, adaptive Branch-Site Random Effects Likelihood (aBSREL)^33^. aBSREL tests whether a given lineage has undergone significant positive selection and estimates the ω ratio, which is the ratio of the non-synonymous to synonymous substitution rates (dN/dS). We found that GBP5 experienced bursts of positive selection during mammalian evolution, in particular in rodents, primates, lagomorphs and bats **(Fig. 2A).** We further analyzed each mammalian order independently, using BUSTED^34^ from HyPhy^35^. We confirmed that mammalian GBP5 has evolved under positive selection in several orders **(Fig. 2B)**, particularly during Rodentia and Chiroptera evolution, as well as in Artiodactyla and Lagomorpha.

### GBP5 is under strong diversifying evolution in bats at the genomic and genetic levels

Because we found evidence of positive selection in bat GBP5 at the gene-wide level, as well as pairwise-amino acid sequence divergence that could reach 20% within Chiroptera **(Fig. S2),** we performed comprehensive evolutionary analyses in this order. To identify when positive selection has occurred during bat evolution, we used aBSREL^33^ from the Chiroptera-specific GBP5 alignment. We found that several bat lineages were the targets of intensive episodic positive selection, including the branch at the origin of the Microchiroptera suborder, as well as other internal and terminal lineages spread-out in the Yangochiroptera and Yinpterochiroptera clades **(Fig. 3A-B).** This indicates differential selective pressure during bat evolution and may represent lineage-specific adaptations driven by past viral epidemics.

**Figure 3.**
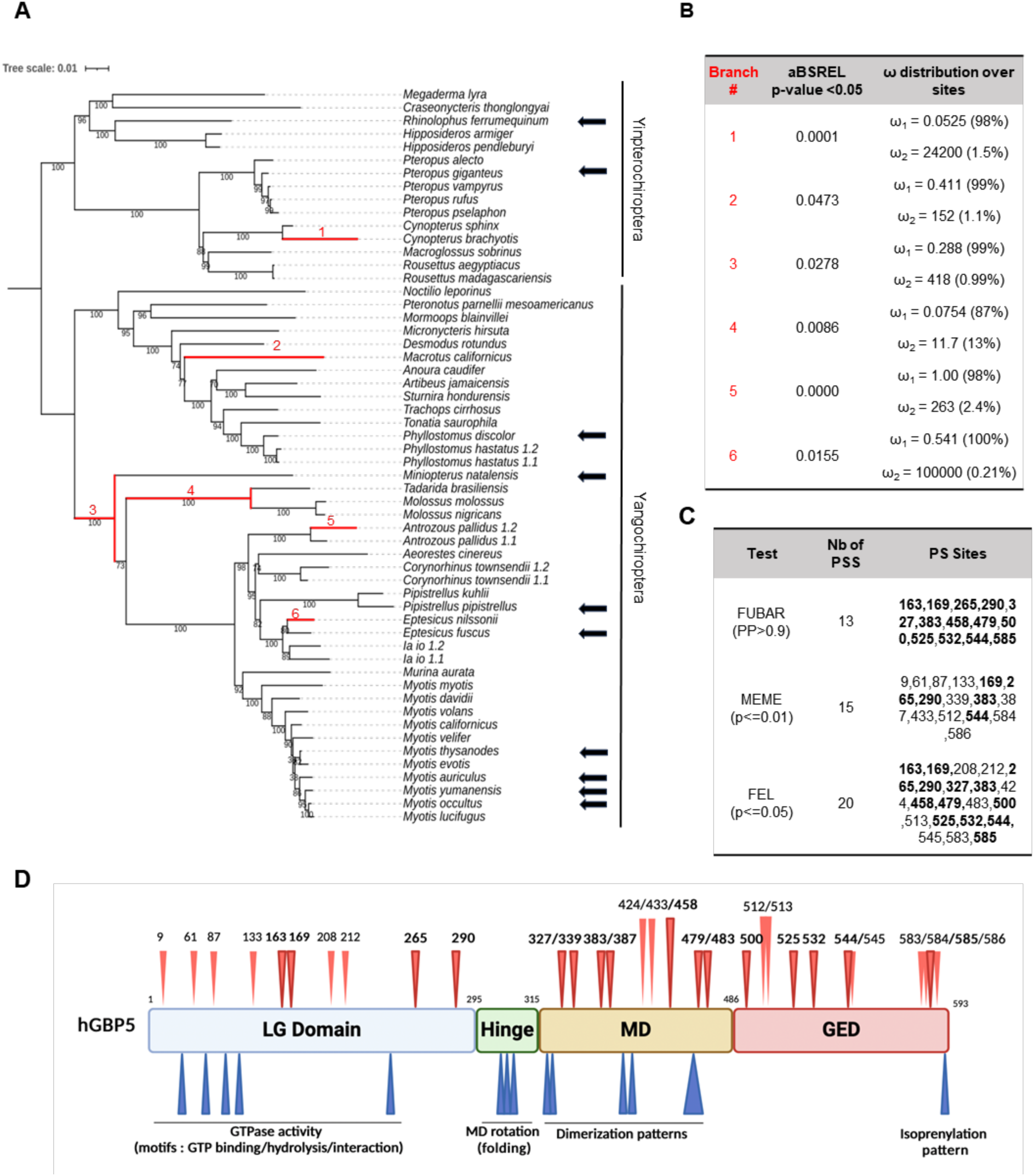
GBP5 has evolved under strong positive selection in bats. A, Phylogenetic and positive selection analyses of bat GBP5. Maximum likelihood phylogenetic tree of bat GBP5 was built with IQ-TREE and statistical support is from 1000 bootstrap replicates (values are shown below branches). Branches under significant positive selection (p-value <0.05) assigned by aBSREL are in red and the corresponding estimated values of ω are reported in panel B. The scale bar indicates the number of substitutions per site. The black arrows identify species with GBP5 genes that were functionally tested in this study. 1.1 and 1.2 identify GBP5 duplicates within a given bat species. B, Evidence of lineage-specific positive selection during bat GBP5 evolution. aBSREL identified at least six branches under significant positive selection. ω1 and ω2, estimation of ω in the rate class not allowing positive selection, and allowing positive selection (ω>1), with % of sites in this class in parentheses, respectively. C, Sites under positive selection (PS) in bat GBP5. Site-specific positive selection analyses were performed using FUBAR^36^, MEME^37^ and FEL^38^ from HYPHY/Datamonkey^35,39^. Only the sites above the indicated “statistically significant cut-off” are shown (PP, posterior probabilities for FUBAR; p-value for MEME and FEL). In bold are the sites identified by several methods. Nb of PSS, number of positively selected sites. “PS sites” numbering is according to the codon numbering in the PRANK codon alignment. D, Schematic representation of GBP5 with its functional domains and the herein identified sites under positive selection (red arrows at the top). LG domain, large GTPase domain. MD, Middle domain. GED, GTPase effector domain. The size of the domains is not to scale. Residues identified by at least two positive selection methods are highlighted in bold. In blue, sites or motifs involved in known functions in human GBP5.

Beyond evidence of lineage-specific positive selection, we identified multiple, independent duplication events in bat GBP5, specifically in *Phyllostomus hastatus, Antrozous pallidus, Corynorhinus townsendii* and *Ia io* **(Fig. 3A).** These Yangochiropteran species belong to different families and are separated by a relatively long evolutionary timescale, suggesting independent convergent evolution **(Fig. 3A).** Despite all the duplication events being recent (i.e., only at terminal branches), we found important divergence between the copies **(Fig. 3A, S2),** suggesting possible neo- or sub-functionalization.

Furthermore, we carried out several analyses to specifically identify the sites that have evolved under positive selection. Using three maximum-likelihood methods from HyPhy (see Methods), we found many positively selected residues in bat GBP5 (**Fig. 3C**). Conservatively, considering only residues identified by at least two methods, we identified 16 positively selected sites – of note, if we consider sites identified by at least one method, we have 29 sites total. The rapidly evolving residues are found in all GBP5 protein domains, except for the hinge domain, which contains no positively selected sites **(Fig. 3D)**. Overall, we identified a very high concentration of sites under positive selection, as well as numerous indels (insertion/deletion) and lineage-specific early stop codons in the C-terminal part of bat GBP5.

Taken together, our transcriptomic and phylogenomic approaches **(Fig. 1-3**) indicate that GBP5 is a key innate immune factor engaged in a molecular conflict in bats. This suggests that GBP5 could play important and lineage-specific roles against various pathogens and that past epidemics may have shaped bat innate immunity.

### Natural variation in subcellular localization of bat GBP5

To test how past genetic diversification may have impacted GBP5 functions, we focused on ten bat species (indicated by the black arrows in **Fig. 3A**) that represent: (i) a large panel of orthologous GBP5 sequences encoding for maximum amino acid variations at the sites identified under positive selection (**Fig. S3**) and (ii) covering various bat families of different clades – some of which are very distant from each other while others are very close (e.g., *Myotis spp*.). This approach allowed us to functionally cover the evolution of bat GBP5 over a period of around 60 million years, at both ancient and more recent times. In humans, the localization of GBP5 at the trans-Golgi network (TGN) is absolutely required for its antiviral activities^16–18^. Its localization is driven by a prenylation sequence present at the C-terminal end of the protein (CaaX motif) that anchors GBP5 at the TGN membrane. Synthetic mutation in this domain prevents both TGN localization and antiviral activities (*Homo sapiens* GBP5-C583A) ^16,40^.

To study the subcellular localization of bat GBP5, we cloned the ten selected orthologous bat GBP5s **(Fig. 3A)** with an HA tag in a eukaryotic expression vector, and compared them to human GBP5 (wt), a prenylation-defective mutant (human GBP5-C583A) and a control vector (i.e., not expressing any GBP5). We ectopically expressed GBP5 in TZM-bl cells and imaged the cells 48 hours later by confocal immunofluorescence microscopy. We found co-localization of GBP5 with the TGN46 marker for 8 out of 10 bat species **(Fig. 4),** showing GBP5 localization to the trans-Golgi network similarly as human GBP5. Surprisingly, two closely related bat species *Eptesicus fuscus* and *Pipistrellus kuhlii* exhibited a diffused subcellular distribution in the cell, similarly to what we observed for the *Homo sapiens*-C583A mutant **(Fig. 4).** Thus, bat GBP5s present natural variations in their subcellular localization, from trans-Golgi network to cellular diffusion.

**Figure 4.**
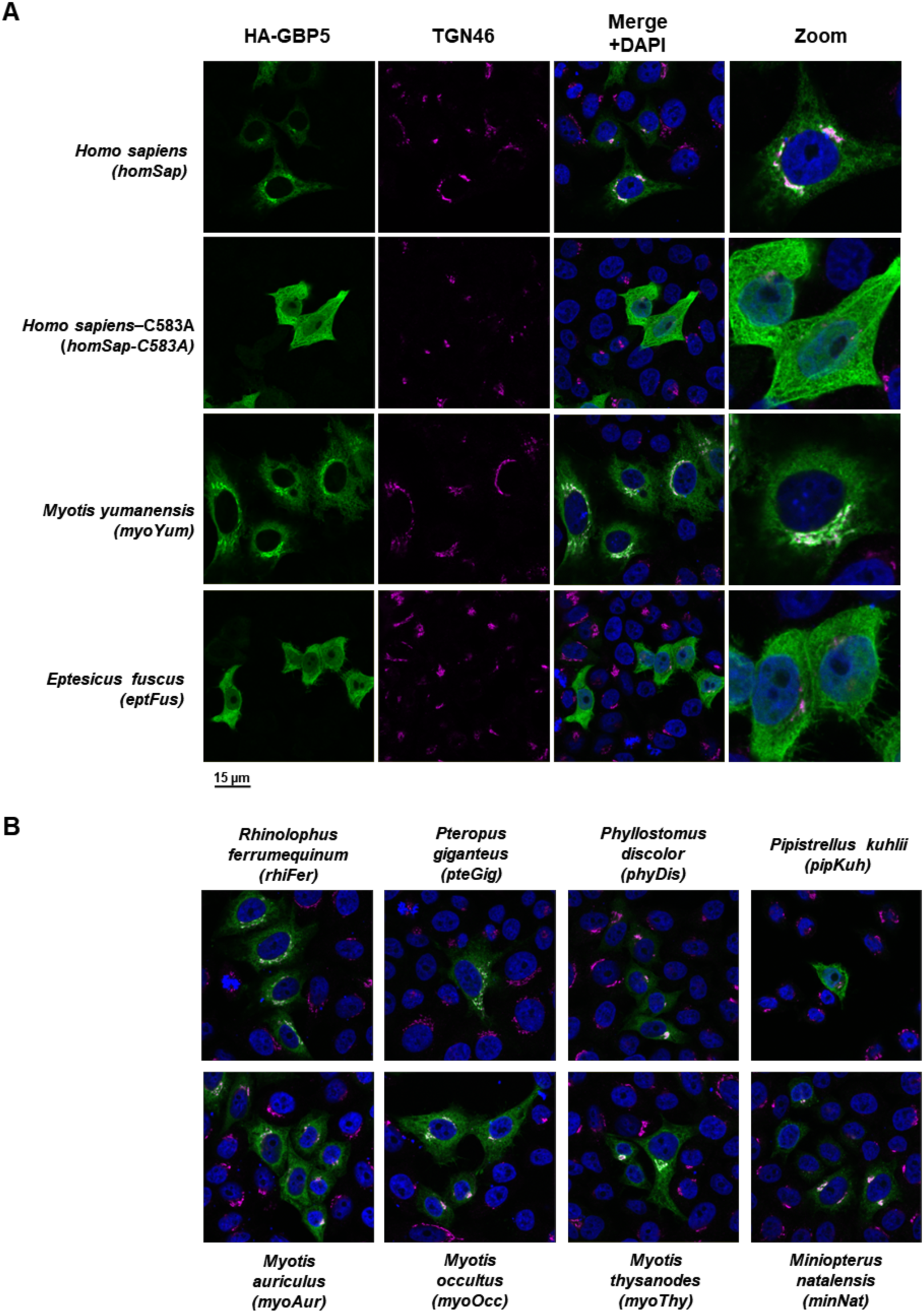
Natural variation in subcellular localization of bat GBP5s. TZM-bl cells were transfected with a plasmid coding for indicated HA-GBP5 species proteins: 10 bat orthologs, and 2 human GBP5s: wt and mutant C583A. Two days post-transfection, GBP5 localization was analyzed by confocal fluorescence microscopy with the indicated marker. Nuclei and trans-golgi-network (TGN) were stained with DAPI and anti-TGN46, respectively. A, All the channels and a zoom are shown for *Homo sapiens*, the mutant *Homo sapiens-C583A, Myotis yumanensis* and *Eptesicus fuscus*. B, Only the merge is shown for the remaining bat species. The complete panel is shown in **Fig. S4**. The pictures present representative results observed in 3 independent experiments. Scale bar indicates 15 μm.

### Bat GBP5s restrict the retrovirus HIV-1 in a species-specific manner, at both viral production and intrinsic infectivity steps

The antiviral functions of human GBP5 have been particularly studied in the context of HIV-1 infection^16,18^, where it was shown to target HIV-1 envelope glycoproteins, reducing the infectivity of neo-synthesized virions. Specifically, GBP5 alters the cleavage of the envelope precursor gp160 in the TGN, preventing maturation into gp120 and gp41 proteins, thereby leading to the production of immature viral particles. However, the precise mechanisms and whether additional inhibitory effects exist remain unclear. Recent reports showed that human GBP5 specifically alters HIV-1 gp120 glycosylation, which can be experimentally witnessed by a typical electrophoretic mobility shift of viral glycoproteins^16,18^. Nonetheless, the anti-retroviral effect of human GBP5 is strictly dependent on its TGN localization.

We therefore assessed the antiviral activity of bat GBP5s against HIV-1 replication on (i) the overall amount of produced virion particles, and (ii) their infectivity, after normalization in virion amount (also called “intrinsic infectivity”). HEK-293T cells were co-transfected with increasing doses of GBP5 or control vector, as well as a plasmid encoding the HIV-1 envelope (Env NL4.3) and an HIV-1 genome plasmid expressing all other HIV-1 proteins and a luciferase reporter (**Fig. 5A** for the experimental setup). Two days later, titration of viral particles in the supernatant, by quantifying viral reverse transcriptase (RT) activity, showed that high expression of bat, as well as human, GBP5 proteins tended to decrease viral particle production, by up to 10-fold reductions for some species (**Fig. 5B**, **Table S1**).

**Figure 5.**
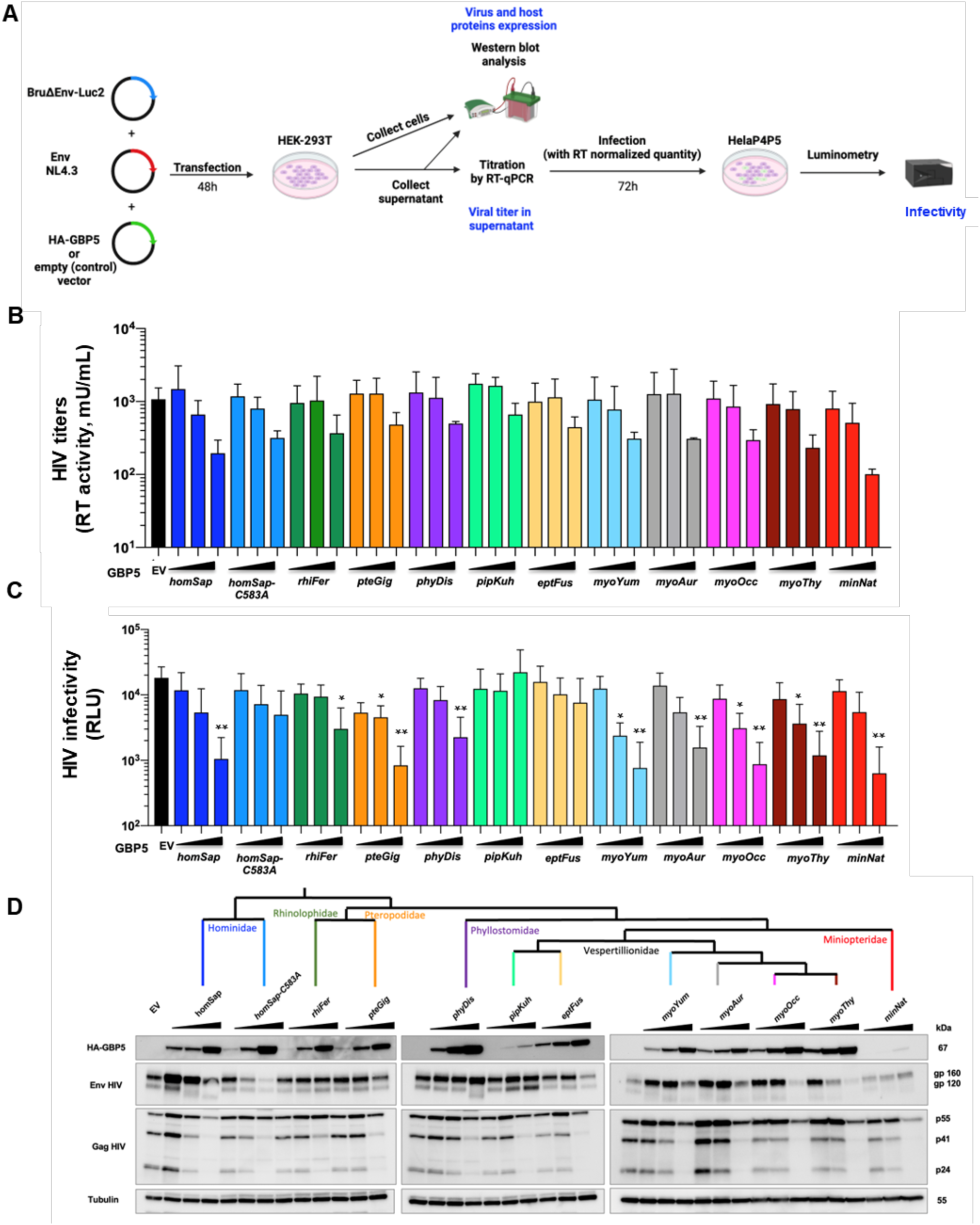
Species-specificity in bat GBP5 restriction of retrovirus (HIV-1). A, Experimental setup. Briefly, HEK-293T cells were transfected with plasmids coding for HA-GBP5 or the empty vector (control), and for HIV-1 LAI genome and Luciferase reporter (Bru∂EnvLuc2 vector), NL4.3 Envelope and Rev. 48h post-transfection, cells were harvested for western-blot analysis, and supernatant for western-blot analysis after ultracentrifugation, titration of virus by RT-qPCR (RT activity), and infection of the RT-normalized viruses in HelaP4P5 cells. Infection was quantified 72h later by luminescence from the viral-encoded Luciferase reporter. B, HIV-1 titers in the supernatants as quantified by RT activity (mU/ml) in the context of a dose of HA-GBP5 (1, 2 or 4 µg) or control vector (EV). The corresponding species of GBP5 is shown (name follows the UCSC nomenclature, three letters from genus followed by three letters from species). C, Corresponding HIV-1 infectivity (RLU, Relative light units). Statistics versus the corresponding control condition: *, p value <0.05, **, p value <0.01. D, Corresponding western blot analysis of HA-GBP5, HIV-1 Env and HIV-1 Gag, and Tubulin (loading control) from the lysates of the HIV-1 producer cells in the context of a GBP5 dose. The cladogram at the top shows the phylogenetic relationships of the tested species. The western blot of the purified virion fraction (supernatant) is in **Fig. S5**.

To determine if bat GBP5 influenced the infectivity of the neosynthesized viral particles, we infected HeLa P4P5 cells (i.e. Hela cells stably expressing the HIV-1 receptors and co-receptors) with a normalized amount of HIV-1 virions (equivalent to 30 mU RT) (**Fig. 5A**). Three days post-infection, we found that several bat GBP5s also efficiently inhibited the virion infectivity of HIV-1 in a dose-dependent manner, similarly to human GBP5 (**Fig. 5C, Table S1**). As examples, *Pteropus giganteus (pteGig), Miniopterus natalentis (minNat), Phyllostomus discolor (phyDis)* and *Myotis spp*. (myoXxx) GBP5s significantly restricted HIV-1 infectivity, with up to 1-log restriction compared to the control condition (EV). However, the GBP5s from three bat species exhibited a distinct behavior: the GBP5s derived from *Rhinolophus ferrumequinum (rhiFer)* showed lower HIV-1 restriction and the *Pipistrellus kuhlii (pipKuh)* and *Eptesicus fuscus (eptFus)* GBP5s presented no infectivity inhibition, similarly to the human GBP5-C583A mutant (**Fig. 5C**). These phenotypic antiviral differences were independent from differences in GBP5 protein expression levels (**Fig. 5D**). Thus, intrinsic species-specific variability in bat GBP5 governs anti-retroviral activity, at the virion infectivity step.

To further characterize virion particles produced in the presence of GBP5s, we analyzed their protein composition by Western blot (**Fig. 5D, S5** for Western blot of virions). We observed that restrictive bat GBP5s reduced virion incorporation of mature gp120 and increased levels of its immature gp160 precursor. This was most often concomitant with a change in the electrophoretic mobility of Env, possibly reflecting an altered glycosylation of gp120 (**Fig. 5D**), as in human GBP5^19^. Yet, we found a case in which GBP5 from *Pteropus giganteus* (pteGig) strongly restricted HIV-1, at both the viral production and the infectivity steps, without any electrophoretic mobility of Env, suggesting a possible different mechanism of antiviral activity that arose in this lineage (**Fig. 5**).

Finally, in the cells and in the supernatant, we observed a generalized GBP5 dose-dependent decrease in the steady-state protein expression of Gag, and possibly Env. This inhibitory effect was found in most GBP5s, including the human C983A mutant, but with varying degrees; for example, *Miniopterus natalensis (minNat)* GBP5 was a strong restrictor with complete Gag p24 expression shutoff in the cell and supernatant (**Fig. 5D, S5**), coherent with the 1-log reduction in the viral titers measured by RT activity (**Fig. 5B, Table S1**). Importantly, we showed that this depletion was not the result of a general cytotoxic effect of GBP5s, because their expressions did not affect cell viability (**Fig. S6**). Thereby, beyond previously described phenotypes and independently of its prenylation, GBP5 may therefore have additional antiviral functions that target other elements or steps in the HIV-1 replication cycle, leading to the inhibition of HIV-1 protein expression and viral particle production.

### Virus-specificity and increased antiviral breadth in bat GBP5s

To determine the breadth of the bat GBP5 antiviral restriction, we tested their effect on the infectivity of viral particles bearing glycoproteins from two other RNA viruses: VSV and EBLV-1. Previous studies showed that human GBP5 inhibits the infectivity of lentiviruses (HIV) pseudotyped with the glycoprotein of EBVL-1 (isolated from *Eptesicus serotinus* bats) but not with that of VSV^18^. Yet, a recent report by Veler *et al.* 2024 suggests that human GBP5 may restrict HIV:VSVg infectivity^19^.

Here, HEK-293T cells were cotransfected with GBP5 or the control vector, and plasmids encoding HIVΔenv luciferase reporter viruses pseudotyped with EBLV-1g or VSVg (HIV:EBLV-1g or HIV:VSVg, respectively). We choose the intermediate dose of 2 µg of GBP5 for these assays, as viral restriction at this concentration was present without toxicity. We quantified viral production from the supernatant, as previously described, and measured the virions’ infectivity in new HEK-293T target cells (**Fig. 6A** for the experimental setup). First, for both pseudoviruses, we similarly observed a decrease in viral production in the supernatant for all GBP5 orthologs (**Fig. S7**), most probably resulting from the restrictive effect of GBP5 on HIV Gag structural proteins (**Fig. 5D, S5**). Second, and after normalization of virion amounts, we found that bat GBP5s reduced the infectivity of HIV:EBLV-1g, but in a species-specific manner in the magnitude of restriction (**Fig. 6B**). GBP5 from *Pteropus giganteus, Phyllostomus discolor, Miniopterus natalensis* and *Myotis spp* significantly reduced HIV:EBLV-1g infectivity by 70-80%. Interestingly, the same inhibitory effect was observed with the human GBP5 and GBP5-C583A mutant, suggesting a restriction mechanism independent of GBP5 isoprenylation at the CaaX motif. In *Rhinolophus ferrumequinum*, *Pipistrellus kuhlii* and *Eptesicus fuscus*, the restriction was milder in the range of 35-45%. By comparison with HIV:HIV-1-Env restriction, these are the same species with the least impacting antiviral effect. This suggests that genetic variations in these bat GBP5s are probably located at common interfaces required for restriction of EBLV-1 and HIV infectivity (i.e., common determinants). Third, we found that most bat GBP5s, as human GBP5, had a little inhibitory effect on HIV:VSVg infectivity. Yet, GBP5 from two *Vespertilionidae* bat species, *Myotis occultus* and *Pipistrellus kuhlii*, significantly restricted HIV:VSVg infectivity (up to 80% inhibition) (**Fig. 6B**). In the latter case, it was particularly unexpected given the lack of HIV infectivity restriction and little protein expression of *Pipistrellus Kuhlii* GBP5. It therefore supports that GBP5 antiviral functions can be virus-specific. It further suggests that specific genetic innovations in GBP5 from some *Vespertilionidae* enabled a gain in antiviral activity, contributing to specificities unique to them. Altogether, these results show that the antiviral restriction of bat GBP5 is species- and virus-specific.

**Figure 6.**
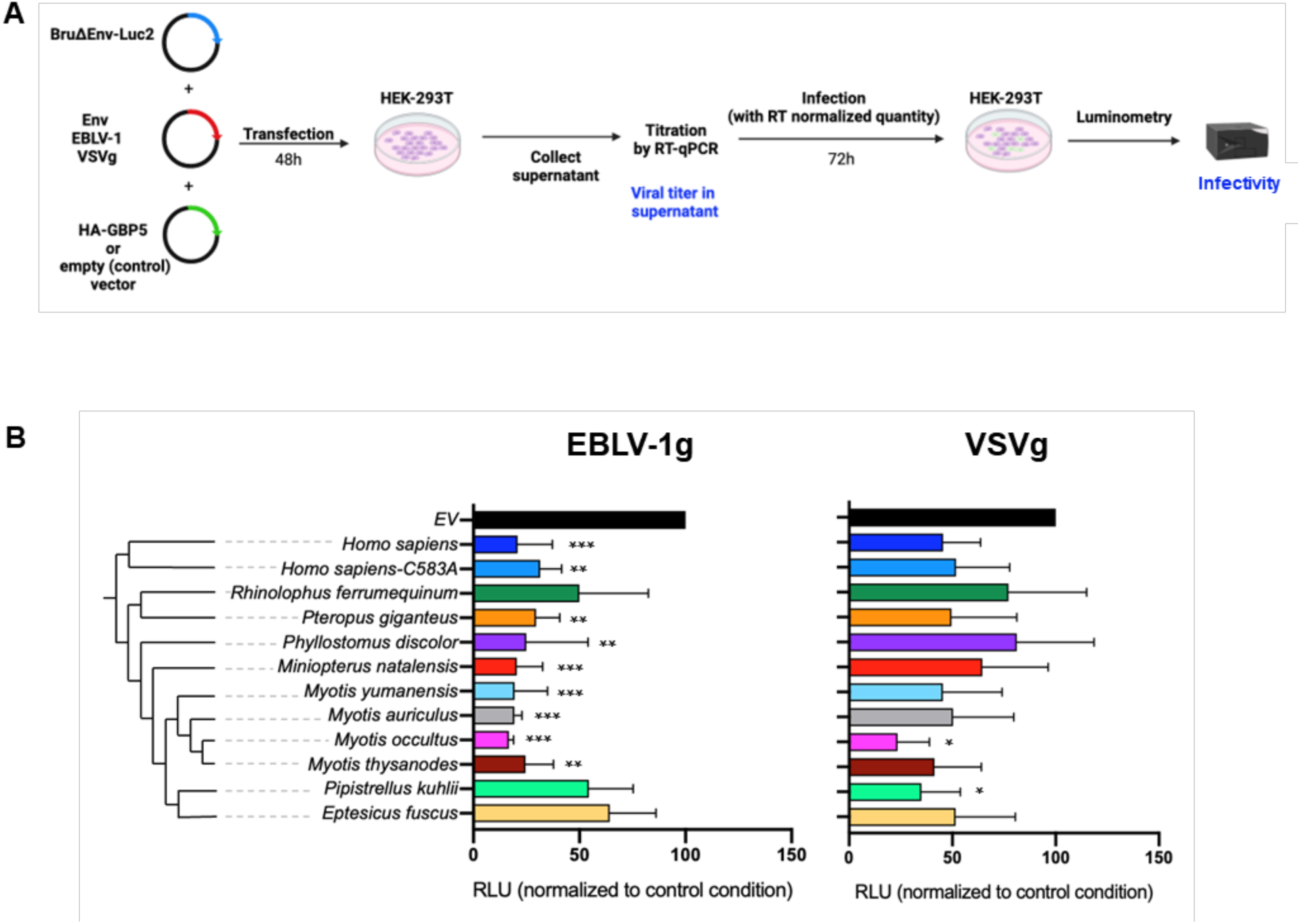
Bat GBP5 restriction is both species- and virus-specific. A, Experimental setup similar to Fig. 5A but with viral pseudotyping with VSVg (VSV condition) or EBLV-1 envelope (EBLV-1 condition). B, Infectivity of RT-normalized pseudotyped-viruses produced in the presence of GBP5, normalized to the condition without GBP5 (EV, Empty vector control) at 100%. The cladogram on the left shows the phylogenetic relationships of the tested GBP5 species. RLU, Relative light units. Statistics versus the corresponding control condition: *, p value <0.05, **, p value <0.01, ***, p value <0,001.

### The CaaX prenylation motif of GBP5 has naturally been lost through early stop codon in one lineage of *Vespertilionidae* bats

To identify molecular determinants of the species-specific subcellular distribution and antiviral activity of bat GBP5s, we performed comparative phenotypic and phylogenetic analyses. Remarkably, we uncovered that several species (n=6) from the same taxon in the *Vespertilionidae* family share an early, fixed stop codon in their genetic sequence (**Fig. 7A, S3**, complete alignment: see Data availability). This includes *Pipistrellus kuhlii* and *Eptesicus fuscus,* which have remarkably different subcellular localization and reduced antiviral functions (**Fig. 4-6**). Importantly, this genomic evidence is strongly supported because it was obtained from publicly available sequences, as well as from *de novo* Sanger sequencing from IFN-induced GBP5 mRNA transcripts of *Eptesicus fuscus* cell lines (see Methods). The acquisition of the early stop codon certainly occurred through a single nucleotide mutation (C to T) in the common ancestor of this clade, approximately 22.6M years ago, leading to a non-synonymous change: “CGA” coding for Arginine (R) to “TGA” Stop codon (**Fig. 7A**). Of note, the nucleotide sequence of the corresponding GBP5 mRNA transcripts, after the early stop codon, did not greatly diverge (**Fig. 7A, S3**, complete alignment available), confirming a relatively recent emergence of this mutation. Interestingly, this natural change led to the complete loss of the prenylation CaaX motif in the GBP5 C-terminal region (**Fig. 7A, S3**, complete alignment available) and this correlated with different subcellular localization and antiviral functions.

**Figure 7.**
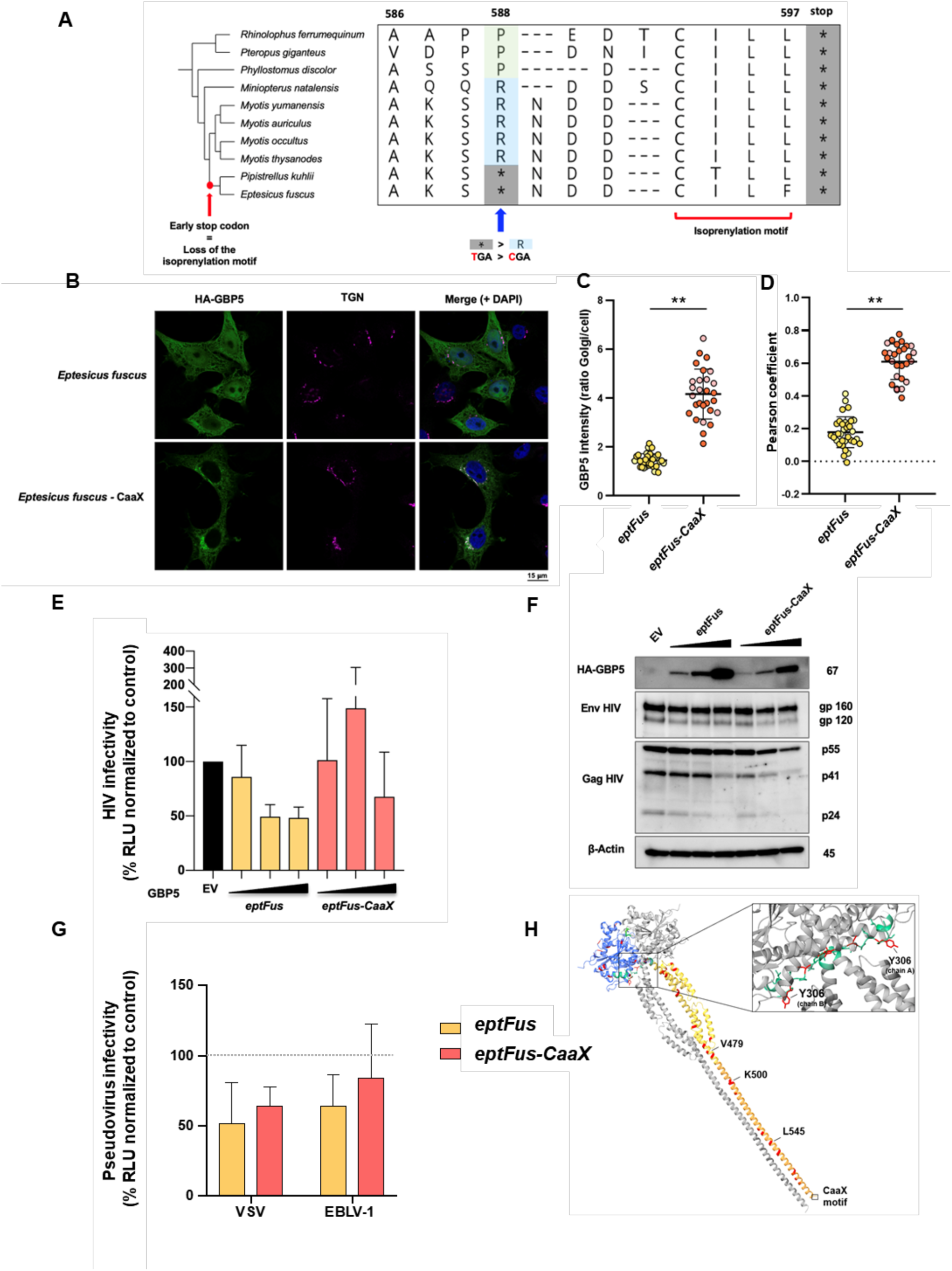
Ancestral reconstruction of the prenylation motif in *Eptesicus fuscus* relocalizes GBP5 to the TGN, but is not sufficient to retrieve full antiviral functions. A, Ancestral state sequence reconstruction upstream of the *Eptesicus fuscus-*CaaX prenylation motif. C-terminal end of the protein alignment of the 10 bat GBP5s tested in functional assays (asterisk, stop codon). Phylogenetic tree was used to infer the ancestral sequence of the C terminal region, the branch where the prenylation motif was lost by a premature stop codon is annotated on the tree. The site of mutagenesis for reconstruction is indicated by the blue arrow. B, Reconstruction of the C-ter relocalizes *Eptesicus fuscus* GBP5-CaaX to the trans-golgi network (TGN). Briefly, TZM-bl cells were transfected with plasmids encoding HA-GBP5s and, 48h later, were analyzed by confocal fluorescence microscopy. Nuclei and TGN were stained with DAPI and anti-TGN46, respectively. Scale bar indicates 15 μm. C, GBP5 mean intensity at the Golgi versus the total cell was quantified for the wild-type *eptFus* and the mutant *eptFus-CaaX*. Each dot corresponds to one cell. Replicates are grouped according to dot color. D, Pearson coefficient correlation per cell calculated between GBP5 and TGN signals for the wild-type *eptFus* and the mutant *eptFus-CaaX.* E-G, Ancestral reconstruction of the prenylation CaaX did not increase *Eptesicus fuscus* GBP5 antiviral functions against infectivity. E, Infectivity of RT-normalized HIV-1 pseudotyped viruses in the presence of GBP5, normalized to the condition without GBP5 (EV, Empty vector control) at 100%. Dose of GBP5 plasmids: 1, 2, 4 µg. Experimental setup as in Fig. 5A. RLU, Relative light units. Viral titers are shown in Fig. S5. F, Corresponding western-blot showing the expression of HA-GBP5, HIV-1 Env and Gag in the viral producer cells with beta-actin as loading control (kDa, on the right). G, Infectivity of RT-normalized VSVg or EBLV-1g pseudotyped viruses in the presence of GBP5 variants, normalized to the condition without GBP5 (empty vector control) at 100%. Experimental setup as in Fig 6A. H, 3D protein structure prediction (AlphaFold) of the reconstructed *Eptesicus fuscus*-*CaaX* GBP5 dimer. Colored and grey chains each correspond to a monomer. Blue, GTPase domain. Green, hinge domain. Yellow, middle domain. Orange, catalytic domain. Red, residues different from *Myotis yumanensis*. Statistics versus the corresponding control condition: **, *p*-value < 0.01.

### Ancestral reconstruction of the prenylation motif in *Eptesicus fuscus* relocalizes GBP5 to the TGN, but is not sufficient to retrieve full antiviral functions

The loss of prenylation signal in another unrelated ISG, the OAS1 (2′-5′-oligoadenylate synthetase 1) protein, was recently described in *Rhinolophidae* bats and was associated with a loss of antiviral activity against SARS-CoV-2^41^. Here, we reconstructed the C-terminal part of GBP5 to rescue its prenylation motif. We used site-directed mutagenesis of *Eptesicus fuscus* GBP5-encoding plasmid to replace the early stop codon with the ancestral amino acid arginine (**Fig. 7A**, *Eptesicus fuscus-*GBP5-CaaX), restoring a long ancestral GBP5. Interestingly, while *Eptesicus fuscus* GBP5 had a cytoplasmic and nucleus diffused distribution (**Fig. 4**), the restoration of the prenylation motif led to its relocalization to the trans-Golgi network (**Fig. 7B, 7C-D** for quantification of immunofluorescence analyses), thereby restoring the ancestral subcellular localization.

Next, we investigated the impact of *Eptesicus fuscus-*GBP5-CaaX on the infectivity of the diverse pseudoviruses previously tested in this study. Surprisingly, the rescue of the CaaX motif was not sufficient to enhance the antiviral effect of *Eptesicus fuscus* GBP5 against the envelope glycoproteins of HIV, EBLV-1 and VSV (**Fig. 7E-G**). Moreover, *Eptesicus fuscus-*GBP5-CaaX, despite being in the TGN, did not induce retention of immature HIV gp160 and had no effect on gp120 glycosylation (**Fig. 7F, S4-S5**). However, the CaaX-reconstructed GBP5 appeared to inhibit Gag and Env total protein expression levels more strongly than the modern eptFus-GBP5, both in the producer cells and in the virions (**Fig. 7F, S5**), without impact on cell viability (**Fig. S6**). Of note, this enhanced restriction in viral production was true even though *Eptesicus fuscus-*GBP5-CaaX protein was less stably expressed (**Fig. 7F, S5**). Altogether, this suggests that, beyond the CaaX prenylation motif, other regions/determinants of *Eptesicus fuscus* GBP5 have evolved and are implicated in antiviral mechanisms.

## Discussion

In this study, we found that bat GBP5 displays signatures of genomic and functional adaptations conferring specificities to bat antiviral innate immunity. Our integrative approach supports a model in which bat GBP5 has been engaged in a molecular conflict with viral pathogens during Chiroptera evolutionary history. Indeed, we found evidence of signatures of positive selection, genomic duplications and natural C-terminal truncation of the protein. And importantly, these genomic adaptations contributed to variations in the subcellular localization of bat GBP5 and to species-specific interactions regarding modern viruses. Because GBP5 was the most differentially expressed gene upon interferon stimulation of *Myotis yumanensis* primary cells, this further supports our primary findings of bat GBP5’s importance in viral infections.

At the genomic and functional level, we found the natural loss of the prenylation motif in GBP5 of six species from the same taxon, associated with a loss of localization to the trans-golgi network (TGN) and reduced antiviral activity. This loss happened in the common ancestor by missense mutation leading to an early stop codon. The loss of prenylation motif in important innate immune factors has previously been reported for the dsRNA sensor OAS1 in the bat *Rhinolophidae* family, although the genetic mechanism of loss was different as the latter was due to an ancestral LTR insertion^41^. In that case, the loss of OAS1 prenylation induced a loss of sensing and restriction against SARS-CoV-2. Because *Rhinolophidae* bats are reservoir species of sarbecoviruses, OAS1 prenylation loss may be a specific adaptation allowing tolerance (i.e. a lesser negative inflammatory response) to a large diversity of coronaviruses circulating in these bats. It is also possible that OAS’s anti-coronavirus function was lost as a consequence of lesser pathogenic coronaviruses circulating in these bat species. Likewise, GBP5 prenylation loss may be an adaptation in response to viral infections specific to this Vesper bat lineage (comprising *Eptesicus* and *Pipistrellus*), which are reservoirs for a diverse range of viruses, including retroviruses and RNA viruses (e.g. rhabdoviruses)^42^.

The ancestral reconstruction of *Eptesicus* GBP5 C-terminal with the prenylation motif modified its subcellular localization but, in contrast to *Rhinolophidae* OAS1^41^, failed to rescue its complete antiviral functions, suggesting that other interfaces of the protein have also diverged. The additional determinants may be amongts the sites that have evolved under strong positive selection. These may result from selective pressures from ancestral viral epidemics, in particular at direct virus-host molecular interfaces. For example, sites under positive selection in bat PKR pointed to molecular interfaces, determinants and conflicts with poxvirus antagonists^8^. In the present work, we found clear species-specificity in heterologous virus-host assays (as reviewed in^43^), further supporting that some positively selected sites may be directly involved in antiviral activity. To start looking for the possible determinants, we compared the genetic sequences of GBP5 *Eptesicus fuscus-*CaaX and *Myotis yumanensis*, which share 94.7% aa identity but have different phenotypes (i.e., *Eptesicus fuscus-*CaaX GBP5 has little antiviral function while *Myotis yumanensis* GBP5 has strong antiviral effects) (**Fig. S2, S8**). We identified several divergent residues, highlighted in red on the 3D predictive structure of *Eptesicus fuscus*-CaaX (**Fig. 7H**), including three of them, V479, K500 and L545, that have evolved under strong positive selection in bats (**Fig. 3B**). Genetic mutations appearing at these sites, potentially resulting from virus-host genetic conflicts, could be key interfaces involved in the GBP5 antiviral mechanisms. Furthermore, an additional site Y306, located in the hinge domain, is essential for the conformational change of human GBP5 (in which an aspartic acid is found at the same position), enabling human GBP5 dimerization and its antiviral activity^40^. This mutation could therefore impair correct folding of the protein and compromise its active state.

Another case of such functional-genetic comparative analysis is possible amongst the four closely-related *Myotis* species, which have different antiviral phenotypes and only 13 divergent sites between them (**Fig. S9**). Indeed, *Myotis occultus* showed a stronger restrictive effect against HIV:VSVg than the other *Myotis* species (≃ 80%, vs. 60% of restriction for *Myotis thysanodes*, for example). Residue K547 is the only residue that differentiates *Myotis occultus* from other functionally tested *Myotis* (also different from human GBP5), suggesting a potentially interesting candidate for future functional analysis. Collectively, comparative analysis with *Myotis yumanensis* revealed several residues that would be interesting to functionally test. Further studies are therefore needed to identify the molecular determinants of bat GBP5 antiviral activity.

Bat GBP5 restricts virion infectivity in a species-specific manner, particularly, cleavage and glycosylation of HIV envelope glycoproteins. Human GBP5 activity is linked to cellular furin-dependent inhibition for HIV glycoprotein cleavage, whereas impaired glycosylation is mediated by a furin-independent mechanism^16,18,19^. Recent investigations on SARS-CoV-2 revealed that human GBP5 may affect the activity of the oligosaccharyltransferase (OST) complex, suggesting that any proteins glycosylated by the OST complex in the ER may be impacted by GBP5^21^. Glycoproteins of HIV, EBLV-1 and VSV could interact with this cellular partner and depend on its activity for their glycosylation. Given the inter-species antiviral specificities, it would be interesting to explore these two mechanisms by furin activity and peptide N-glycosidase (PGNase) assays. Furthermore, although the GTPase activity of human GBP5 appeared dispensable to its described antiviral activities^16^, investigating the involvement of bat GBP5 GTPase activity in antiviral activity could reveal new features. Because we observed a repression of HIV Gag structural protein expression in a CaaX motif-independent manner, it is possible that the enzymatic activity of GBP5 may be required for complementary antiviral activities. Furthermore, some bat GBP5s seem to restrict more viruses than human GBP5. It would therefore be interesting to test other viruses and to determine whether bat GBP5s have acquired an extended antiviral breadth and by which mechanism. Given bat diversification and GBP duplications, GBP5 from bats may also have evolved a yet unknown restriction mechanism. Exploring functional impact of natural variations throughout mammalian evolution also contributes to the broad mechanistic understanding of innate immunity.

Beyond the antiviral effector functions, GBP5 also targets a broad spectrum of bacteria and parasites by promoting their eradication via activation of the inflammasome, notably leading to pyroptosis of the infected cell ^14,15,44^. Variations at the positively selected sites could also affect these GBP5 functions. Furthermore, rapidly evolving residues in bat innate immune factors have also been involved in increased antiviral defense, or in beneficial viral tolerance. For example, bat IRF3 positive selection at specific serine residue participated in enhancing antiviral defenses ^45^, or rapid evolution of S358 in bat STING participated in increasing viral tolerance ^46^, respectively. Previous studies also reported that altered expression and function of bat inflammasome components, leading to reduced inflammasome activation, may participate to traits associated with asymptomatic viral reservoirs and greater longevity ^3,47^. Some bat GBP5 adaptations could therefore also be implicated in dampened inflammatory responses, providing interesting insights for understanding viral tolerance mechanisms.

Overall, this study contributes to a better understanding of bat innate immunity, which has been a prime objective. Past pathogenic viral exposures at different times led to modern antiviral immune specificities. This study also demonstrates the importance of including multiple related species in comparative functional studies to better assess divergence in their immune systems, thereby participating to fundamental knowledge of mammalian innate immunity and ultimately contributing to translational research for One Health and therapy^48^.

## Materials and Methods

### Bat sampling

*Myotis spp. (yumanensis, occultus, auriculus and thysanodes)* bats sampled for this study were wild caught under scientific collection permits for California. Bats were sampled using standard mist-netting procedures, including taking standard body measurements, following USGS recommendations for White-Nose Syndrome and COVID-19 prevention. Two 3-mm wing punch biopsies were taken from the left and right plagiopatagium of each donor individual and placed in a live cell collection media consisting of DMEM/F12 (Gibco) supplemented with 15mM HEPES (Gibco), 20% FBS (Gibco), and 0.2% puromycin (Invivogen) ^49–51^. Wing punches were then brought back to a cell culture facility in Berkeley, where they were used to generate cell lines as previously described^49–51^.

### Cell culture

Human embryonic kidney 293T (ATCC, cat. CRL-3216), HelaP4P5 (Hela cells expressing CD4^hi^CCR5^hi^CXCR4^hi^), and TZM-bl (NIH AIDS Research and Reference Reagent Program, Cat. 8129; Hela cells expressing CD4^hi^CCR5^hi^CXCR4^hi^ and the luciferase reporter under an LTR promoter) cells were grown in DMEM supplemented with 5% fetal bovine serum (FCS, Sigma cat. F7524) and 100 U/ml of penicillin/streptomycin (Sigma-Aldrich). Bat primary cells were maintained in DMEM supplemented with 10% fetal bovine serum (FCS, Sigma cat. F7524) and 100 U/ml of penicillin/streptomycin (Sigma-Aldrich).

### RNA extraction for next-generation sequencing and quality assessment

*Myotis yumanensis* primary cells from the three individuals were plated in 12-well plates. Cells from each individual constituted a replicate. Twenty-four hours later, cells were either treated with universal type I interferon treatment (PBL assay science) at 1000 U/mL or the medium was replaced with IFN-free DMEM for the mock condition. After six hours, cells were collected and pelleted, and then stored at −20°C. RNA was extracted from frozen cell pellets according to RNA isolation Macherey Nagel kit protocol. To maximize RNA extraction quality, DNAse incubation time was increased to 30 minutes. Total RNA concentration was quantified using Qubit RNA high sensibility kit and Qubit4 machine (Thermofisher). RNA integrity was assessed by capillary electrophoresis on Tapestation 4150 (Agilent). All samples reached a RNA integrity number superior to 9 and were used for library preparation.

### Library preparation and RNA sequencing (RNAseq)

Samples were enriched in mRNA using NEBNext® Poly(A) mRNA Magnetic Isolation Module (New England Biolabs, E7490-lot n°10151189) following supplier protocol. Enriched mRNA samples were stored at −80°C prior to library concentration. RNA libraries were generated using CORALL Total RNA-Seq Library Prep kit (UDI 12 nt set B1, Lexogen). Libraries control assessment was performed again using Qubit dsDNA HS Assay Kit to perform fluorimetry dosage on Qubit4 and measure libraries profile by capillary electrophoresis Tapestation 4150. After quality assessment, libraries were mixed in equimolar concentration according to Illumina. The sequencing was performed on all samples using three successive runs on Illumina NextSeq 500 plateform in paired-end 78-pb with dual indexing at IGFL sequencing platform.

### Bioinformatic analyses of RNAseq data

#### Data quality assessment

Raw data quality was assessed using FASTQC (version 0.12.0) and results were compiled using MultiQC (version v1.14). The data were trimmed to remove adapters/UMI index and low quality reads using FASTP (version 0.23.1, with the parameter -min_len 40 and -q 20). *Mapping and quantification*.Transcriptome mapping was performed using a transcriptome index (Salmon 1.10.2), based on *M. yumanensis* reference transcriptome supplied by our collaborator from Berkeley and Arizona Universities based on accession number. As the quasi-mapping transcriptome rate, obtained ranged between 40-55% using Salmon, the read were mapped on *M. yumanensis* reference genome using STAR (2.7.11b) to check for potential contamination. The genomic mapping rate had a range between 70-90%. A manually inspection determined this difference of mapping rate is due to the non annotation of UTR in the transcriptome of these non model species. *Table counts and DESeq2 analysis.* Salmon quantification files were imported using tximport (version 1. 26.1) with the option “lengthScaledTPM” and “countFromAboundance”. The gene orthology relationship with *Homo sapiens* was obtained using Orthofinder (version 2.3.10) including 4 others myotis species. This analysis is restricted to genes defined as one-to-one orthologs with Homo sapiens. Differentially expressed genes were obtained using DESeq2 (v 1.42.1) package. The following additional package were used to generate the different figures: ggrepel (version 0.9.5), ggplot2 (version 3.5.1), tidyverse (version 2.0.0) and MetBrewer (version 0.2.0).

#### Comparative analyses with Shaw et al 2017 dataset

Our RNAseq results for GBP5 were compared with results from Shaw et al 2017 (EBI, project accession number PRJEB21332). The cells used for the Shaw *and al* dataset were also primary fibroblast cell lines (cows, 4 individuals; sheep, 3 individuals; dog, 1 individual; human and rodent, PromoCell and European Collection of Authenticated Cell Cultures: C-12302 and 06090769, respectively; *M. lucifugus*, from Oregon, USA)^22^. For the IFN treatment, cell lines were similarly treated with 1,000 U/ml universal type I IFN for 6h, except for dog cell lines treated with 200 ng/ml canine IFNα.

### *De novo* sequencing of GBP5 genes

Total genomic RNA was extracted from bat primary fibroblast cells of *Myotis auriculus, Myotis occultus, Myotis thysanodes,* and *Myotis yumanensis*, as well as from *Eptesicus fuscus* immortalized skin fibroblasts (a gift from Rachel Cosby and Cédric Feschotte^52^) using Macherey-Nagel NucleoSpin RNA kit following the manufacturer’s protocol. Total RNA was reverse transcribed into complementary DNA (cDNA) with random primers and oligo(dT), using the SuperScript III One-Step reverse transcription polymerase chain reaction (PCR) kit (Thermo Fisher Scientific, Poland). GBP5 mRNA was then amplified from each species using 10 ng of cDNA and different sets of primers (Table S2) that were specifically designed using an alignment of publicly available GBP5 sequences. The PCR reactions were performed using the New England Biolabs (NEB) Q5 High-Fidelity DNA Polymerase, following the manufacturer’s protocol, including a final volume of 50 µl, a 0.5 µM primer concentration, and an annealing temperature of 61°C. PCR products with multiple bands were excised and purified from gel using the NucleoSpin Gel and PCR Clean-up Kit from Macherey-Nagel. Sanger sequencing of GBP5 was performed by a commercial company (Genewiz, Azenta Life Sciences, Germany).

### Collection of GBP5 orthologous sequences

Homologs of all six human GBPs were searched in 536 mammalian reference genomes (downloaded from NCBI in June 2023) by sequence similarity using the miniprot software ^53^. Homologous matches that were at least 50% of the length of the human GBP5 were selected. Because genome sequences can include indel errors that destroy the reading frame, a first round of multiple sequence alignment of the retrieved GBP homologs was run with MACSE v2 ^54^. MACSE uses a statistical model that takes frameshifts into account in order to properly align homologous codons. The resulting coding sequences were then processed with PREQUAL^55^ to eliminate segments with insufficient homology and likely to be misaligned. Then, the remaining coding sequences were aligned with MACSE again. The GBP5 orthologs to the human GBP5 were then identified by generating a phylogenetic tree with IQ-TREE2 (under a GTR+R6 model)^29^.

In total, GBP5 coding sequences from Chiroptera (n=55; including 5 newly generated sequences), Primates (n=37), Rodentia (n=80), Artiodactyla (n=97), Carnivora (n=51), Lagomorpha (n=10), Perissodactyla (n=9), Proboscidea (n=3), Pilosa (n=1), Cingulata (n= 4) and Xenarthra (n=1) were retrieved.

### Phylogenetic and positive selection analyses of GBP5 orthologous sequences

GBP5 orthologous codon sequences were aligned for each mammalian group separately using PRANK ^28^ or Muscle ^56^, and were manually curated. A phylogenetic tree was built using the maximum likelihood method implemented in IQ-TREE^29^. Node statistical support was computed through 1000 bootstrap replicates. The gene codon alignments were then submitted to positive selection analyses using the HYPHY package (either through command line or the DataMonkey webserver,^35,39^). First, Branch-site Unrestricted Statistical Test for Episodic Diversification (BUSTED,^34^) was used to determine whether the gene has experienced episodic positive selection (p value <0.05). Second, an adaptive branch-site REL test for episodic diversification (aBSREL)^33^ was used to identify the branches under positive selection with a cutoff at p value <0.1 as well as to quantify the dN/dS ratio for each branch independently. Lastly, we used three statistical analyses to identify the specific sites under positive selection: MEME which detect individual sites subject to episodic diversifying selection (^37^, cutoff p value < 0.01), FUBAR (A Fast, Unconstrained Bayesian AppRoximation for Inferring Selection,^36^; cut off posterior probabilities PP > 0.90) and FEL (Fixed Effects Likelihood, ^38^; cutoff p value < 0.05).

### Plasmids

GBP5 were amplified from bat cDNA (species: *Myotis auriculus, Myotis occultus, Myotis thysanodes, Myotis yumanensis*, *Eptesicus fuscus)* of fibroblast cells (details above) or synthesized by a commercial company (Genewiz, Azenta Life Sciences, Germany) (species: *Rhinolophus ferrumequineum, Pteropus giganteus, Pipistrellus kuhlii, Miniopterus natalensis, Phyllostomus discolor*). They were cloned (*via* BstBI and XhoI restriction sites) into the vector MT06 (RRL.sin.cPPT.CMV/-BamHI-HA-BstBI-E2-crimson-XhoI.IRES-puro.WPRE, shared by Caroline Goujon: Addgene plasmid # 139448^57^), where an HA tag was added between BamHI and XhoI restriction sites. Ancestral reconstruction of the *Eptesicus fuscus* isoprenylation motif expression was generated from wild-type *Eptesicus fuscus* pMT06-GBP5 by replacing the Stop codon with corresponding ancestral residue, using the Quik Change Lightning Site-Directed Mutagenesis Kit (Agilent) following the manufacturer’s instructions (set primers available in **Table S2**). For pseudovirus production, a plasmid coding for LAI-ΔEnv-Luc2 genome (Bru-ΔEnv-Luc2 that has no Env and encodes a firefly luciferase in place of the nef gene, shared by Michael Emerman) was used and pseudotyped with HIV-1 NL4.3 Env (along an HIV-1 Rev plasmid to facilitate nuclear export of Env, ^58^), VSVg (pMD2.G, Didier Trono: Addgene plasmid # 12259) or European bat lyssavirus EBLV-1 Env (pLTR-EBLV1env, a gift from Daniel Sauter^18^).

### HIV-1 production and infection assay

A total of 4×10^5^ HEK-293T cells were seeded in 6-well plates and transfected the next day with plasmids coding for LAI-ΔEnv:Luc2 genome (1.2 µg), NL4.3 envelope (120 ng), HIV-1 Rev (75 ng), as well as indicated doses of HA-GBP5 (0, 1, 2, 4 µg) or empty vector control using HBS/CaCl2 to produce infectious HIV-1 single-round viruses. Forty-eight hours later, cells were harvested for western blot and virus-containing supernatants were clarified for virus titration, by measuring HIV-1 RT activity (see below), western blot (see below) and infections. For infection, 1×10^4^ HeLaP4P5 were seeded into white-clear bottom 96-well plates. Twenty-four hours later, cells were infected with 30 mU RT of virus supernatant in the presence of DEAE-Dextran solution (20 µg/ml). Luciferase activity from the reporter HIV-1: Luc2 virus was measured seventy-two hours post-infection by Tecan Spark® Luminometer.

### VSV and EBLV-1 pseudovirus productions and infection assays

A total of 4×10^5^ HEK-293T cells were seeded in 6-well plates and transfected the next day with Bru-ΔEnv-Luc2 (shared by Michael Emerman) (1.2 µg), VSV-g or EBLV-1 envelope, as well as indicated doses of HA-GBP5 or empty vector control using HBS/CaCl2 to produce infectious pseudotyped virus. Virus-containing supernatant was harvested, clarified for viral titration, by measuring RT activity and infection. For infection, 1×10^4^ HEK-293T were seeded into white-clear bottom 96-well plates and infected with 50 mU RT of virus supernatant. Luciferase activity from the reporter pseudovirus was measured seventy-two hours post-infection by Tecan Spark® Luminometer.

### Virus titration by RT activity

Titration of HIV-1, VSVg and EBLV-1 Env pseudoviruses from supernatant were quantified by measuring reverse transcriptase (RT) activity as in ^59^. Briefly, virions were lysed in a homemade lysis buffer (0.25% Triton X-100, 50 mM KCl, 100 mM Tris−HCl pH 7.4, 40% glycerol) and 0.4 U/μL RiboLock RNase inhibitor (Thermo Scientific). The viral lysates was then diluted in H2O and added to the RT-qPCR mix: 35 nM bacteriophage MS2 RNA (Roche) as a template for RT, 100 nM of each primer (5’-TCCTGCTCAACTTCCTGTCGAG-3’ and 5’-CACAGGTCAAACCTCCTAGGAATG-3’) and Takyon™ No ROX SYBR 2X MasterMix blue dTTP buffer (Eurogentec) in a total reaction volume of 20 μL. Purified RT enzyme (Abnova) was used to make the standard curve. The RT−qPCR reaction was carried out in a Bio-Rad CFX96 cycler with the following parameters: 42 °C for 20 min, 95 °C for 5 min and 40 cycles (95 °C for 5 s, 60 °C for 30 s and 72 °C for 15 s).

### Viability measurement

A total of 1×10^4^ HEK-293T cells were seeded in white clear-bottom 96-well plates and transfected the following day with plasmids encoding the LAI-ΔEnv:Luc2 genome (30 ng), NL4.3 envelope (3 ng), HIV-1 Rev (1.9 ng), as well as a plasmid encoding HA-GBP5 (100 ng) or an empty vector control using TransIT-LT1 (Mirus). Of note, the quantities were calculated to maintain the same ratio used for HIV production with the higher dose (4 µg) of GBP5 described above. Forty-eight hours later, cell viability was assessed by measuring adenosine triphosphate (ATP) levels using the CellTiter-Glo® Luminescent Cell Viability Assay (Promega). Treatment with etoposide (100 µM) for 24h was used as a positive control (Sigma). The luminescence of ATP activity was quantified using a Tecan Spark® luminometer.

### Western Blot and antibodies

Cells were lysed by ice-cold RIPA buffer (50 mM Tris pH8, 150 mM NaCl, 2 mM EDTA, 0.5% NP40) supplemented with protease inhibitor cocktail (Roche) and by sonication. Cell-free virions were concentrated and purified from the supernatant by centrifugation (2h, 30 000 rpm, 4°C) through a 20% sucrose cushion and resuspended for lysis in Exo-RT lysis buffer. Proteins from cell or virus lysates were mixed in Laemmli buffer, heated at 95°C for 5min and separated by SDS-PAGE on 4–20% Bis-Tris gels (Invitrogen) before being transferred to PVDF membrane by an overnight wet transfer at 4°C. Membranes were blocked in a TBS-T 1X solution (« Tris Buffer Saline », Tris HCl 50 mM pH8, NaCl 30 mM, 0.05% of Tween 20) with 5% powder milk. Proteins were stained with a mouse anti-p24 HIV-1 CA (1:1000, NIH HIV Reagent Program, cat.183-H12-5C), a mouse anti-HIV-1 Env (16H3) (1:1000, NIH-ARP, cat. 12559), a rabbit anti-HA (1:5000, Sigma, cat. H6908), a mouse anti-tubulin (1:5000, Sigma, cat. T5168) or a mouse anti-β-actin (1:5000, Sigma, cat. A2228), and then with a donkey anti-rabbit-HRP (1:5000, Sigma, cat.AP188P) or a donkey anti-mouse-HRP secondary antibody (1:5000, Sigma, cat.AP16017). Detection was made with SuperSignal West Pico Chemiluminescent Substrate (ThermoFisher Scientific) using the Chemidoc Imagina System (Biorad).

### Immunofluorescence imaging

A total of 3×10^4^ TZM-bl cells were seeded on glass coverslips and were transfected with 1 µg of GBP5 plasmid with jetPRIME, according to the manufacturers’ instructions (Polyplus). Forty-eight hours later, cells were fixed with 4% paraformaldehyde (PFA) (Sigma) for 15 minutes at room temperature and permeabilized in 0.25% Triton-T×100 (Sigma) for 5 minutes. A blocking step was carried out for 1h at room temperature with 3% BSA (Sigma) and 0.1% Triton T×100 in PBS. Primary antibody incubation was carried out for 1h at RT with rabbit anti-HA (1:500, Sigma, cat. H6908) and sheep anti-TGN46 (1:500, Biorad, cat.AHP500GT) to label respectively HA-GBP5 proteins and the Trans-Golgi-Network (TGN). Primary antibodies were detected with secondary donkey anti-rabbit AlexaFluor-594 (Invitrogen, cat.A212027) and donkey anti-sheep AlexaFluor-647 (Thermofisher, cat.A-21448), for 1h at RT. All cells were labeled with DAPI (4′,6-diamidino-2-phenylindole) (Thermo Scientific) - containing solution (1:10,000 dilution in PBS). Images were acquired using Zeiss LSM 880 AiryScan confocal microscope and analyzed with Imaris software. For quantification analyses, segmentation of the cells was first performed using Cellpose algorithm (version 3.0.11)^60^. Proportion of GBP5 at the TGN and colocalization analysis were then performed on ImageJ software using an automatized macro or the BIOP JACoP plugin, respectively.

### Other software and statistical analyses

Differences between conditions were statistically analyzed using GraphPad Prism 9 software. A Nested t test was used for immunofluorescence analysis, and an ANOVA test for all other data sets. For each of these tests, the p-value was considered significant when inferior to 0.05. Error bars in graphics are SEM or SD (see Figure legends). Schematic workflow figures were created using BioRender.

## Data and Reagent availability statement

All data are publicly available. The scripts are available in gitlab https://gitbio.ens-lyon.fr/ciri/lp2l/gbp5_paper_2025. The RNAseq data were deposited in NIH Bioproject database under accession number PRJNA1011374. The five *de novo* GBP5 sequences obtained for five bat species were deposited in GenBank (GenBank Submission SUB15056411). The alignments for phylogenetic analyses are all openly available in FigShare Dataset (https://doi.org/10.6084/m9.figshare.26180785.v1, https://doi.org/10.6084/m9.figshare.26180764.v1, https://doi.org/10.6084/m9.figshare.26180803.v1, https://doi.org/10.6084/m9.figshare.26180761.v1, https://doi.org/10.6084/m9.figshare.26180767.v1, https://doi.org/10.6084/m9.figshare.26180776.v1, https://doi.org/10.6084/m9.figshare.26180794.v1, https://doi.org/10.6084/m9.figshare.26180797.v1). Reagents are accessible upon request to the corresponding author.

## Acknowledgements

We thank the members of the LP2L team (CIRI) and Molly Ohainle (UC Berkeley) for helpful discussion. We thank Veronika Krchlikova for helpful feedback on the manuscript. We thank Cédric Feschotte and Rachel Cosby (Cornell University, NY) for generously sharing *Eptesicus fuscus* cell lines. We thank Ryan Byrnes and Kathleen Slocum, who assisted with the collection of two of the *Myotis yumanensis* primary cell lines. We thank Molly Ohainle, Michael Emerman and the Emerman Lab members (Fred Hutch) for help in setting up the RT titration assay and sharing reagents. We thank Caroline Goujon (IRIM Montpellier) and Daniel Sauter (Univ. Tubingen) for kindly sharing reagents with us. We thank Benjamin Gillet and Sandrine Hughes from the IGFL platform for the library preparation and the RNA sequencing. We acknowledge the contribution of the SFR Biosciences (UAR3444/CNRS, US8/Inserm, ENS de Lyon, UCBL) ANIRA cytometry and PLATIM platforms, especially Véronique Barateau and Jacques Brocard for their assistance, as well as Didier Decimo of the BSL-3 ENS-Lyon. We thank the IFB, *Institut Francais de Bioinformatique*, for enabling bioinformatic analyses on the IFB servers, as well as the High Performance Computing (HPC) resources supported by the University of Arizona TRIF, UITS, and Research, Innovation, and Impact (RII) and maintained by the UArizona Research Technologies department. Finally, we thank all the contributors of the publicly available bioinformatic programs and genomic sequences.

## Author contributions (CRediT: Contributor Roles Taxonomy)

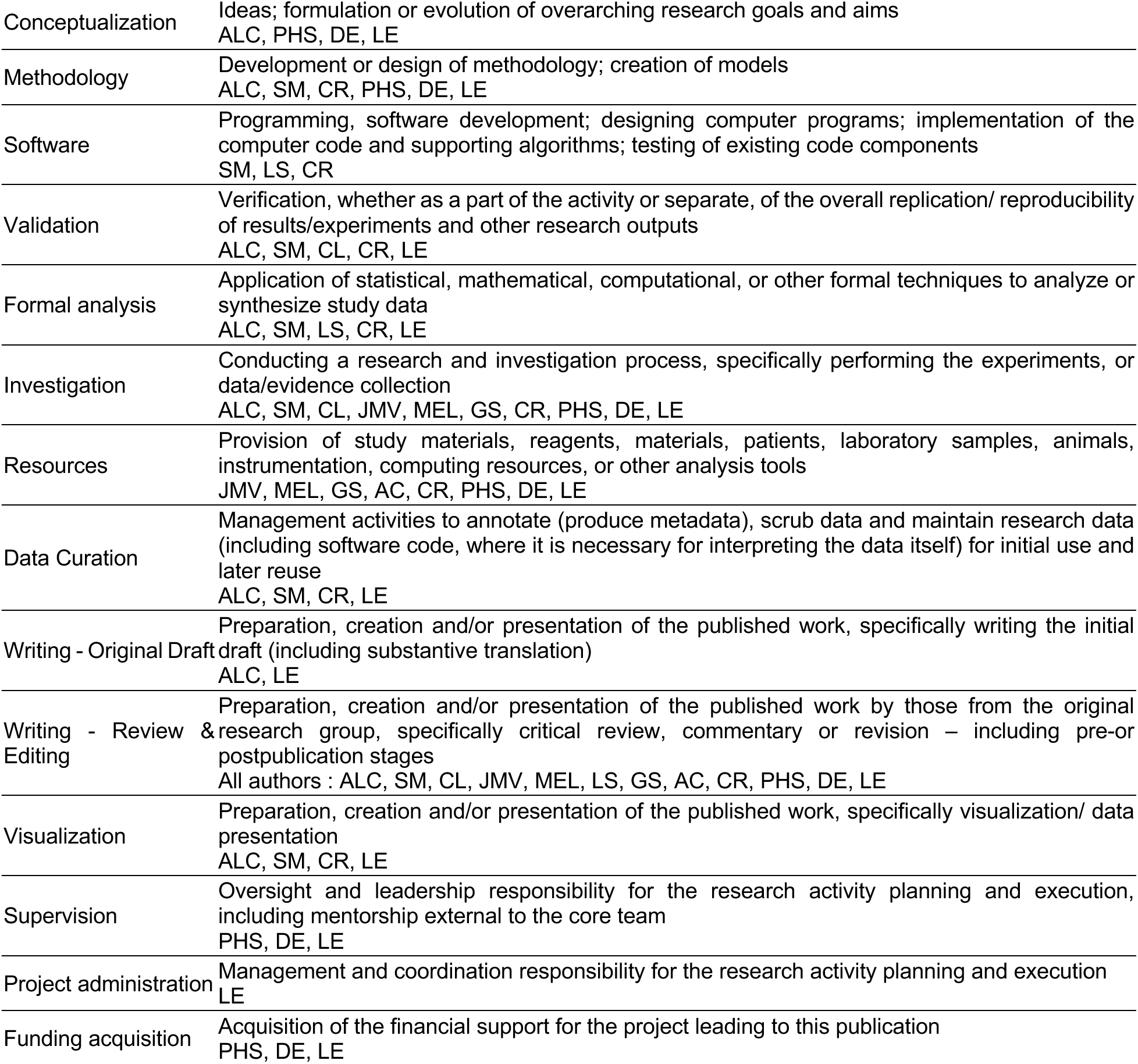

## Supplementary Figures and Tables

**Figure S1 :**
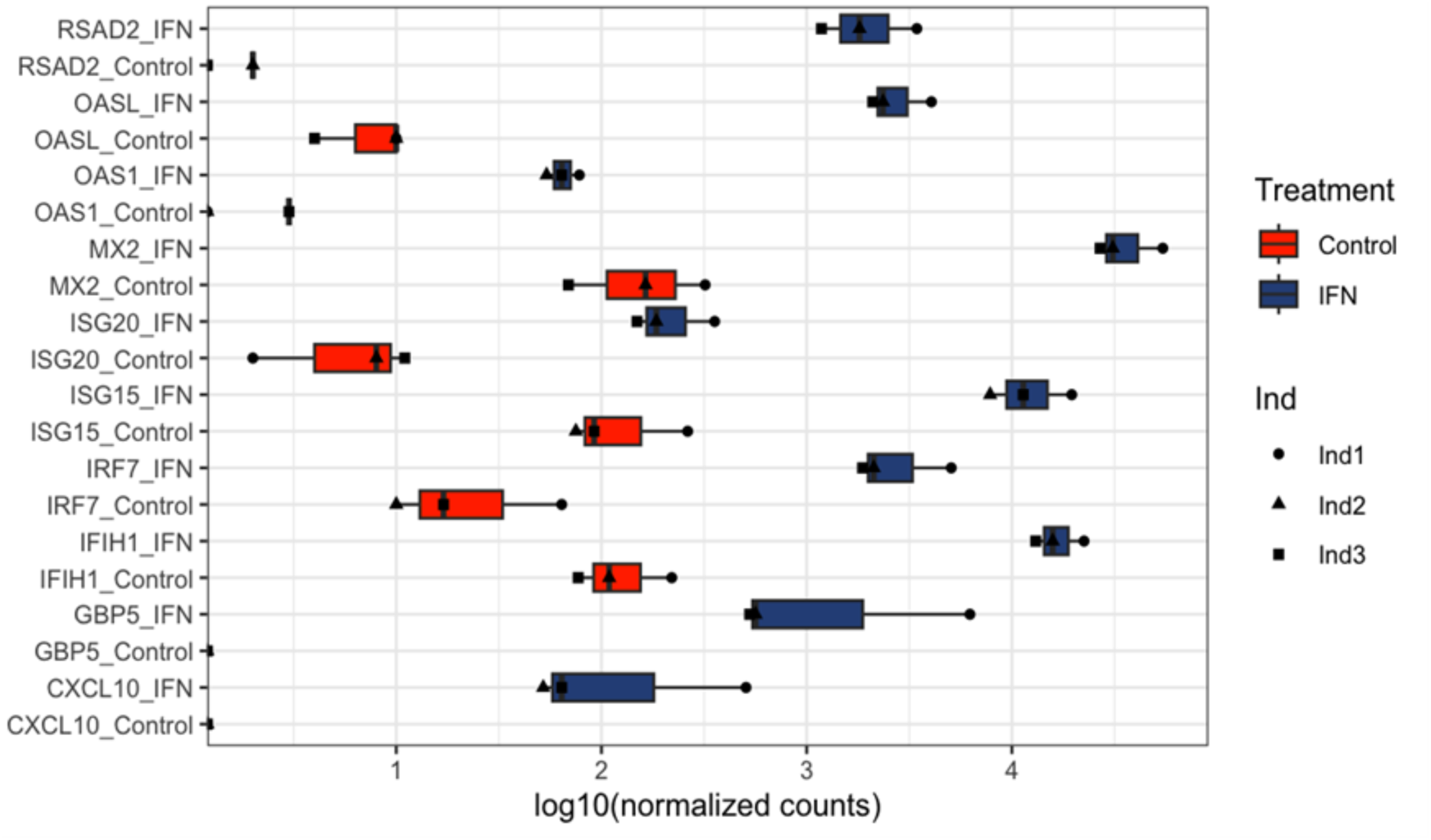
Expression of known ISGs in *Myotis yumanensis* primary cells. Normalized counts in control (red) and universal type I IFN treatment (dark blue) for three individuals (Ind1-3).

**Figure S2.**
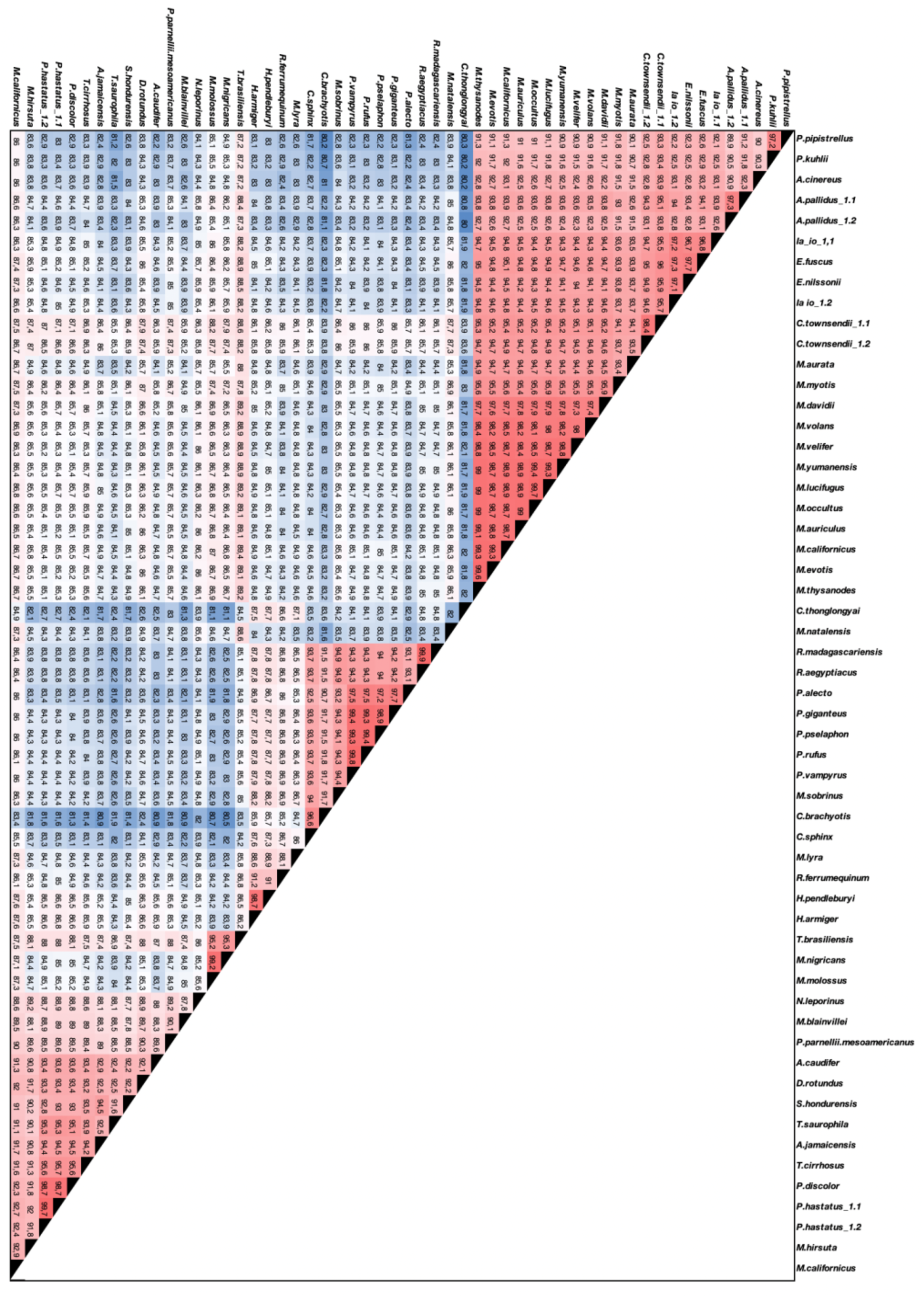
Nucleotide sequence divergence of bat GBP5s. Bat GBP5 sequences were aligned with MUSCLE and percentage of identity measured with GeneiousR10.

**Figure S3.**
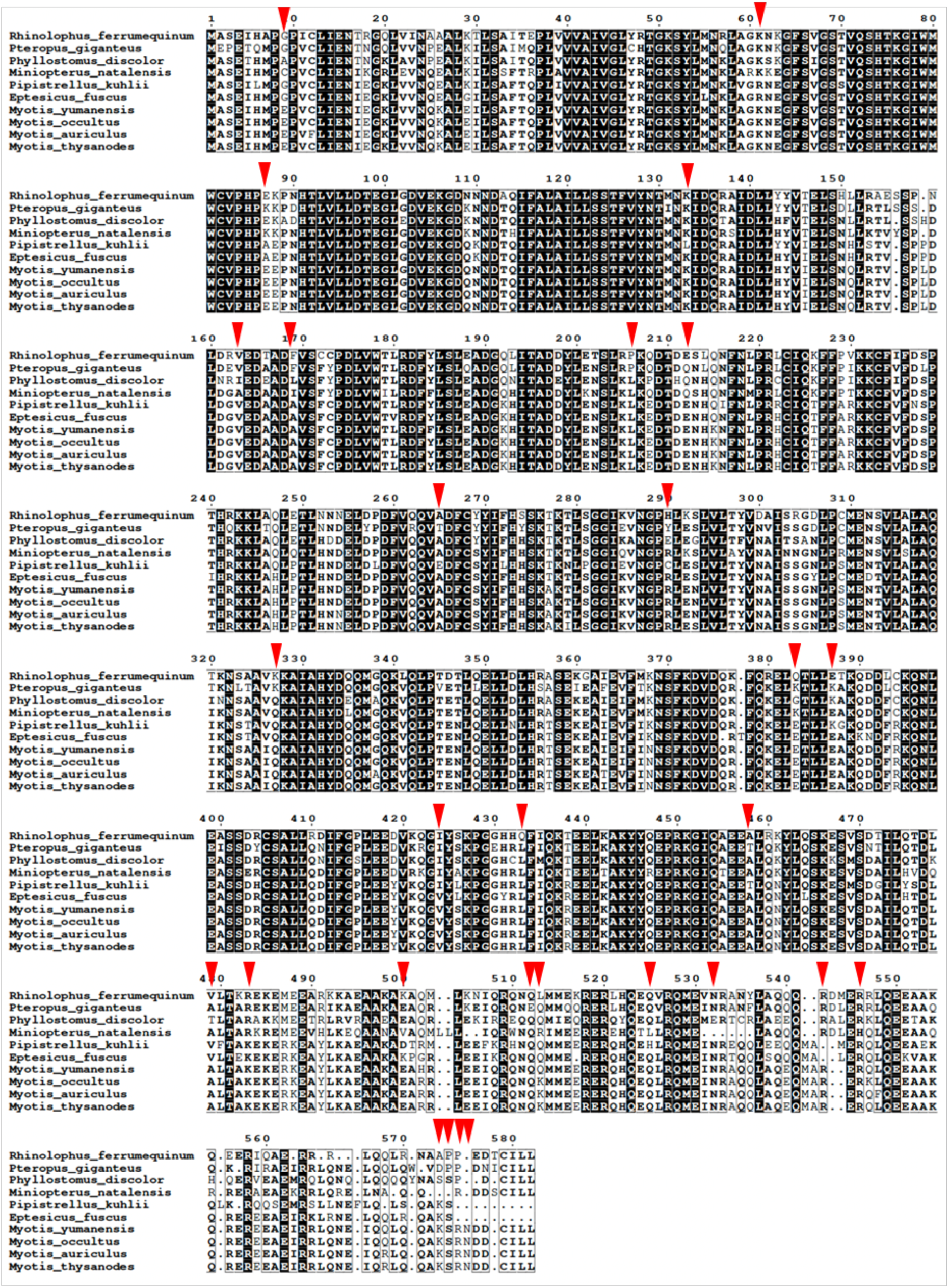
Alignment of functionally tested bat GBP5s. Alignment of functionally tested amino acid sequences from the bat GBP5. In red, positively selected sites in Figure 3C.

**Figure S4.**
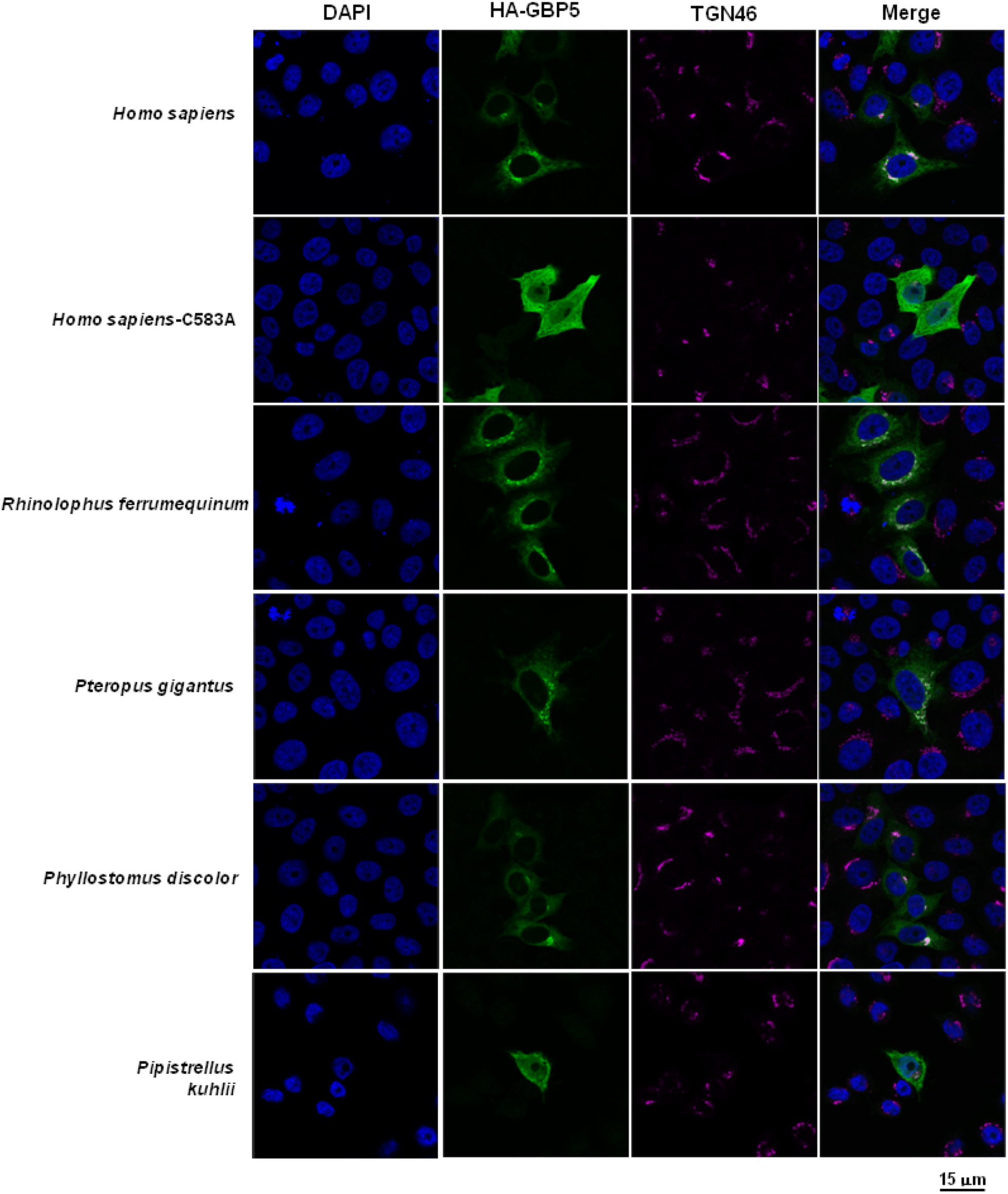
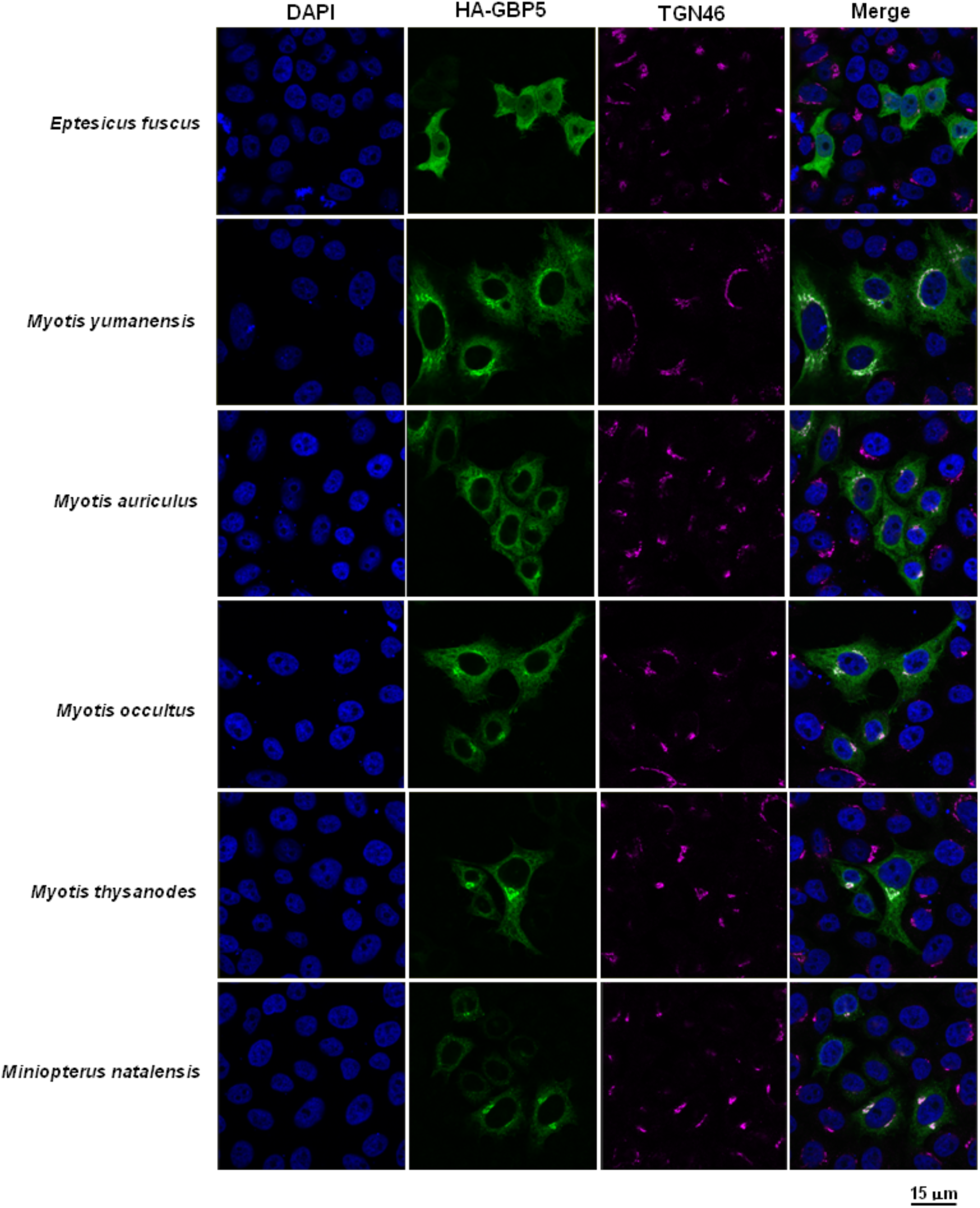
Natural variation in subcellular localization of bat GBP5. TZM-bl cells were transfected with a plasmid coding for indicated HA-GBP5 species proteins. Two days post-transfection, GBP5 localization was analyzed by confocal fluorescence microscopy. Nuclei and trans-golgi-network (TGN) were stained with DAPI and anti-TGN46, respectively. Scale bar indicates 15 μm.

**Figure S5.**
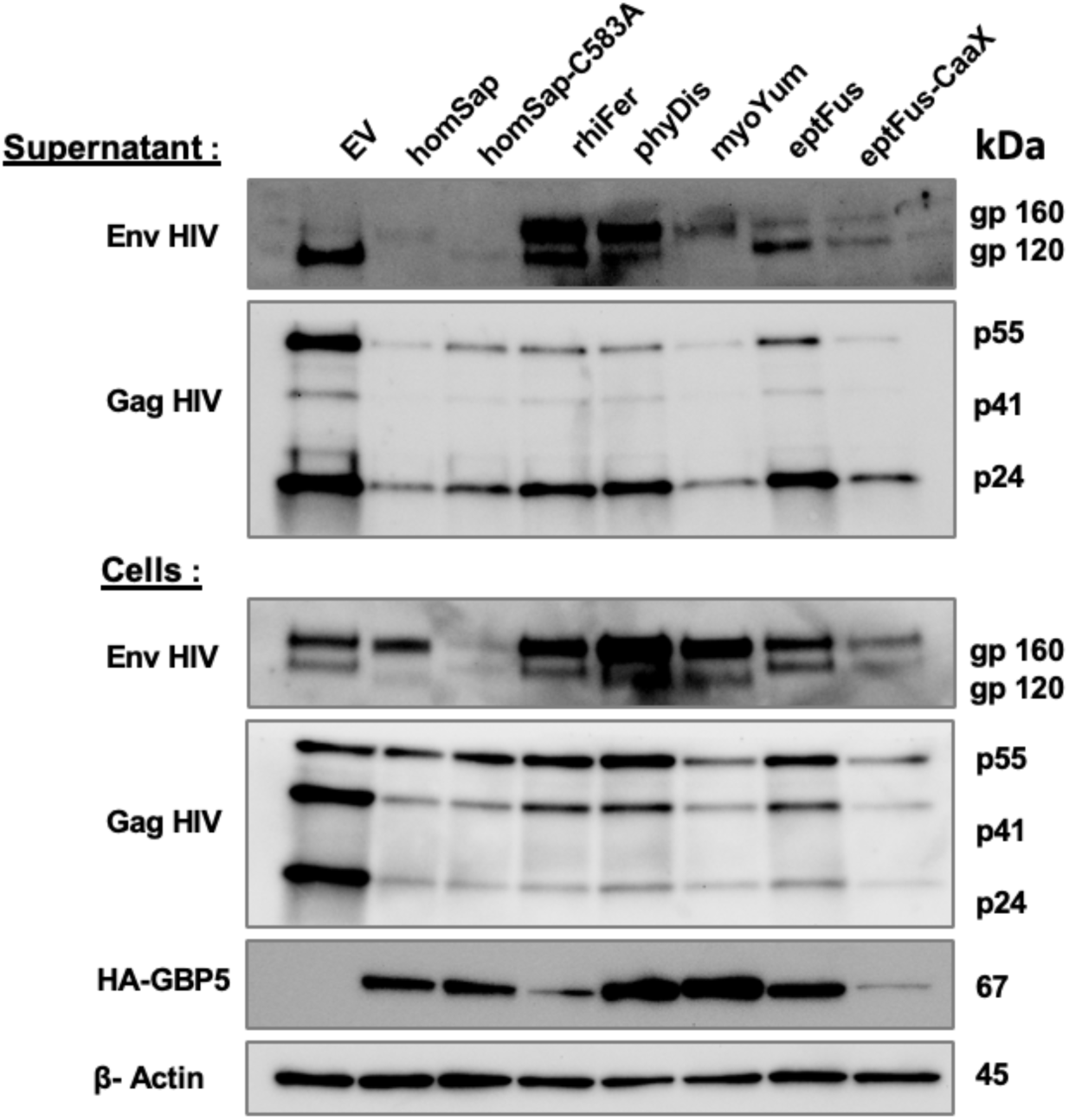
Species-specific restriction of bat GBP5 on HIV-1 Env glycoprotein maturation and viral protein expression. Western blot analysis of HA-GBP5, HIV-1 Env and HIV-1 Gag, and beta-actin (loading control) from the lysates of the HIV-1 producer cells (bottom) and the purified virion fraction of the supernatant (top) in the context of 4µg of the corresponding GBP5 or control vector (EV).

**Figure S6.**
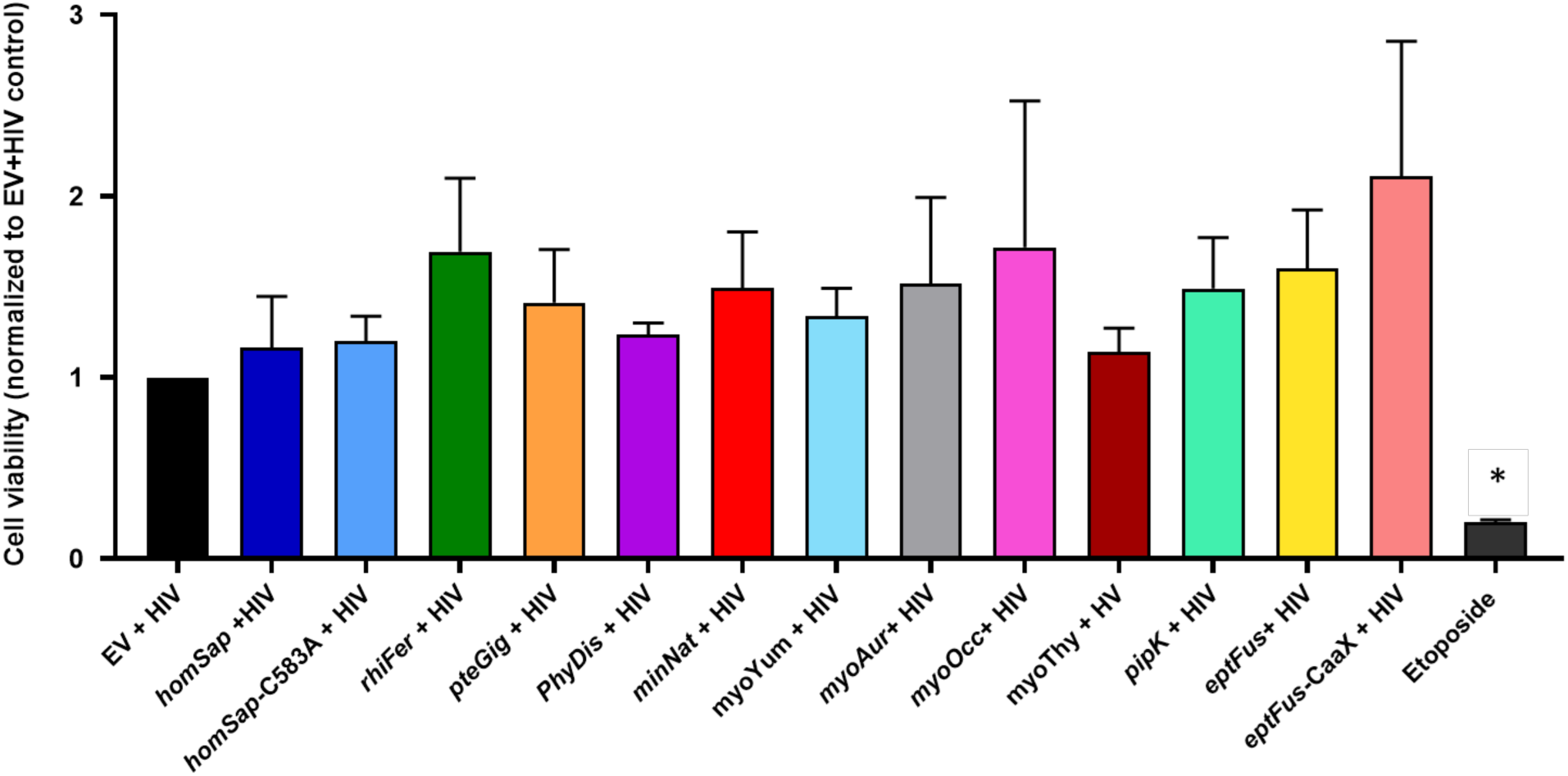
Expression of bat GBP5 in the context of pseudoviral particle production does not increase cytotoxicity in HEK-293T cells. HEK-293T cells were transfected with plasmids coding for HA-GBP5 or the empty vector (EV control), and for HIV-1 LAI genome and Luciferase reporter (Bru∂EnvLuc2 vector), NL4.3 Envelope and Rev (identical conditions used for HIV infection in Fig. 5). 48h post-transfection, cell viability was determined by measuring the level of adenosine triphosphate (ATP). A treatment with etoposide (100µM) during 24h was used as a positive control. Mean values of three independent experiments are shown. Statistics versus the corresponding control condition: *, *p* value <0.05.

**Figure S7.**
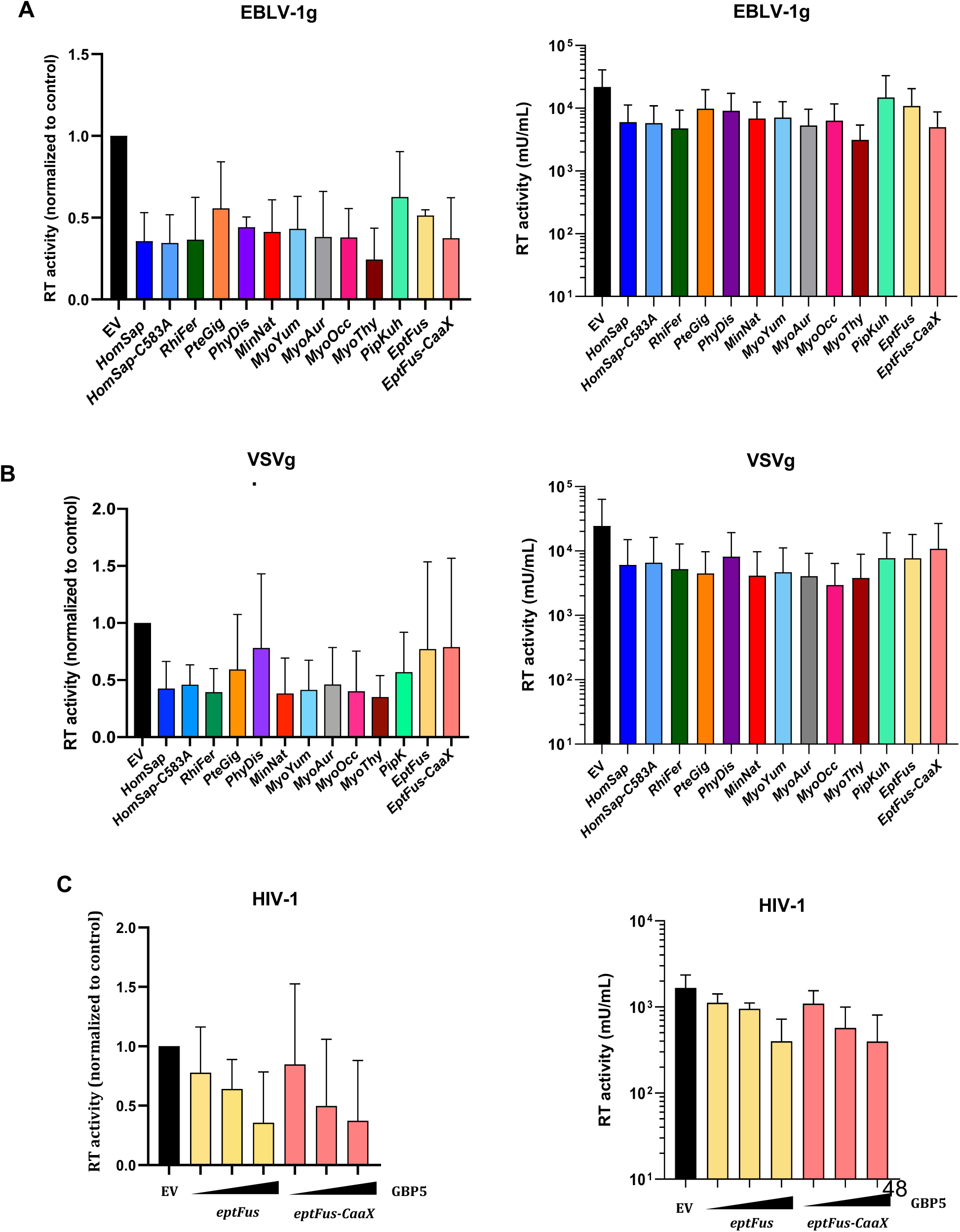
Titration of pseudoviruses by RT activity in the supernatant. Titration of lentiviruses pseudotyped with EBLV-1g (A), VSVg (B), or HIV-1 Env (C) from supernatants, quantified by RT activity (mU/ml). A-B, with 2 µg of HA-GBP5 or control vector (EV) for the corresponding indicated species (i.e. HomSap, Homo sapiens). C, in the context of a dose of HA-GBP5 (1, 2 or 4 µg) or control vector (EV). EptFus-CaaX corresponds to the C-ter ancestral reconstructed *Eptesicus fuscus* GBP5 bearing the CaaX prenylation motif. Left, Normalized values to control (EV) at 1. Right, Raw data (mU/mL).

**Figure S8.**
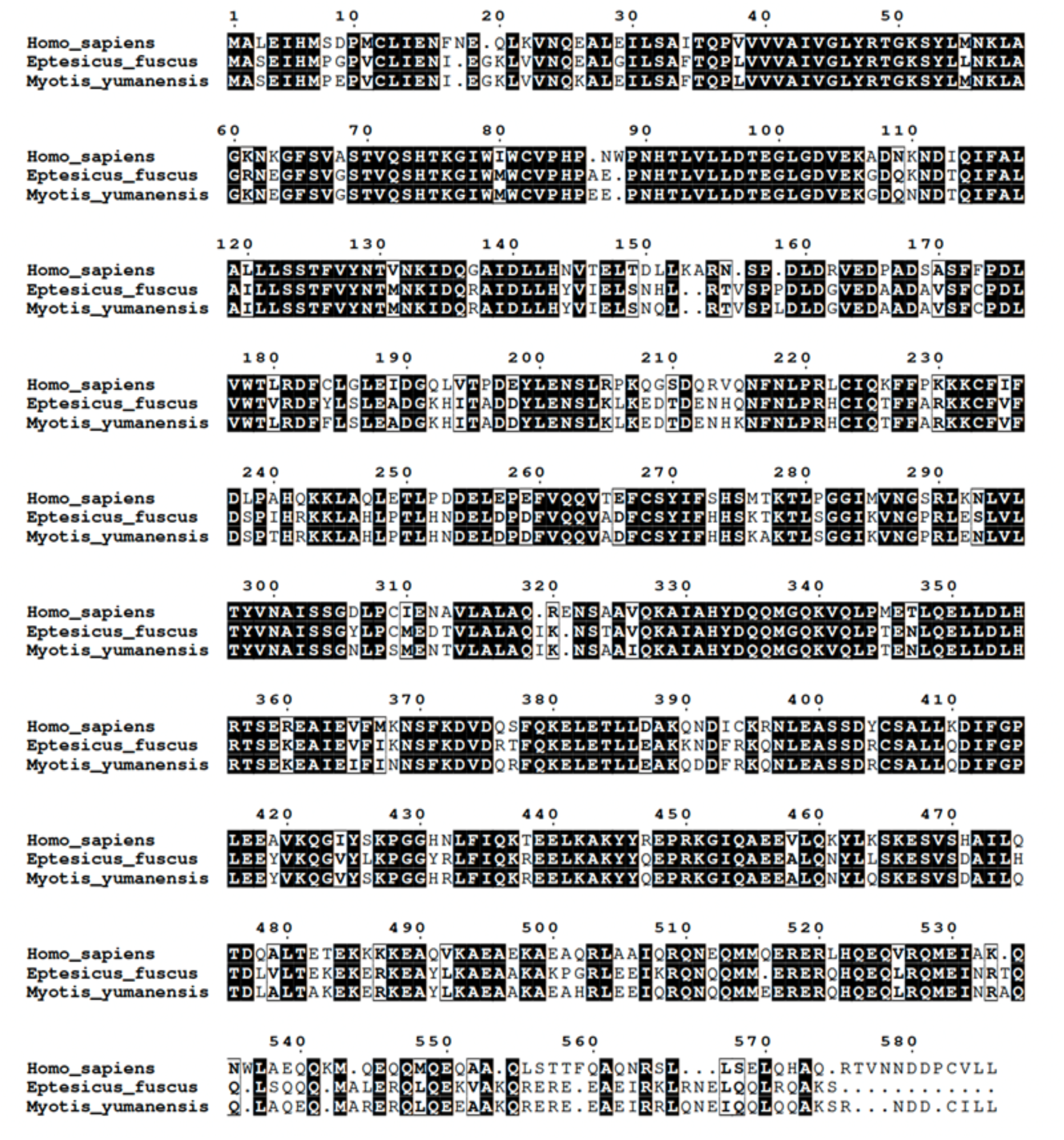
Alignment of *Myotis yumanensis, Eptesicus fuscus* and *Homo sapiens* GBP5 protein sequences.

**Figure S9.**
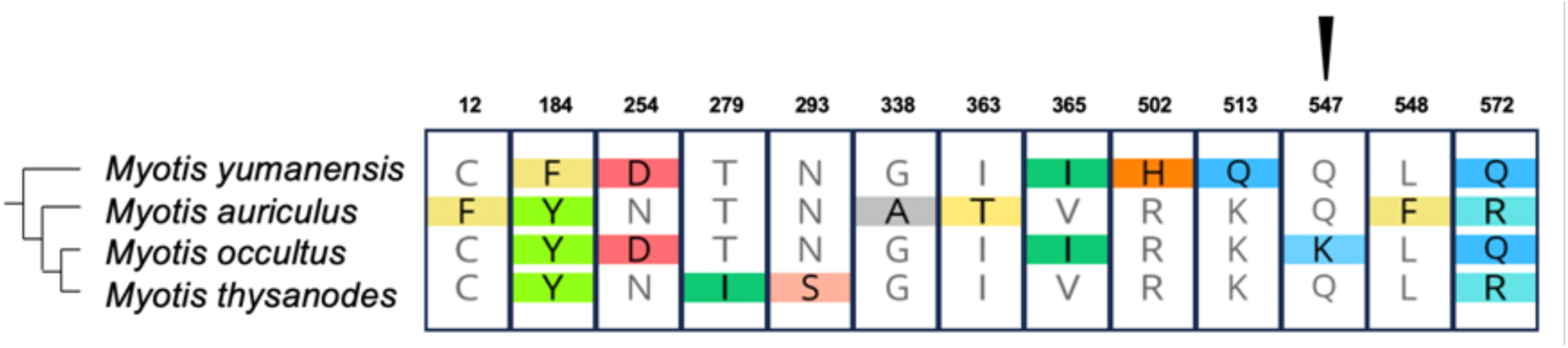
Divergent residues identified between functionally tested *Myotis* GBP5s. Of note, GBP5 from *Myotis occultus* was a stronger restrictor of the infectivity of viral particles bearing VSVg, as compared to other *Myotis* tested (Fig. 6).

**Table S1.** Bat GBP5s inhibit both HIV-1 intrinsic infectivity and viral production, in a species-specific and dose-dependent manner. Summary of the effects of bat GBP5s on the HIV-1 intrinsic infectivity step and on the viral production step.

| Species | GBP5 (ug) | % HIV-1 intrinsic infectivity | % viral production |
| --- | --- | --- | --- |
| EV | 0 | 100,0 | 100,0 |
| homSap | 1 | 56,2 | 119,4 |
|  | 2 | 23,7 | 61,9 |
|  | 4 | 5,6 | 21,8 |
| homSap-C583A | 1 | 62,0 | 109,9 |
|  | 2 | 35,8 | 75,9 |
|  | 4 | 23,4 | 33,2 |
| myoAur | 1 | 73,8 | 106,0 |
|  | 2 | 27,2 | 97,2 |
|  | 4 | 8,2 | 33,4 |
| myoOcc | 1 | 61,7 | 101,7 |
|  | 2 | 17,3 | 72,8 |
|  | 4 | 4,5 | 35,7 |
| myoThy | 1 | 48,8 | 80,4 |
|  | 2 | 19,4 | 70,1 |
|  | 4 | 5,6 | 28,9 |
| myoYum | 1 | 69,1 | 87,6 |
|  | 2 | 12,7 | 63,5 |
|  | 4 | 3,3 | 34,8 |
| pteGig | 1 | 29,9 | 120,8 |
|  | 2 | 34,0 | 129,6 |
|  | 4 | 4,6 | 59,4 |
| pipKuh | 1 | 60,6 | 166,3 |
|  | 2 | 60,3 | 161,8 |
|  | 4 | 111,5 | 75,5 |
| minNat | 1 | 63,6 | 72,0 |
|  | 2 | 25,1 | 44,0 |
|  | 4 | 2,5 | 11,5 |
| eptFus | 1 | 82,8 | 85,7 |
|  | 2 | 54,7 | 98,0 |
|  | 4 | 34,4 | 52,0 |
| rhiFer | 1 | 60,0 | 91,7 |
|  | 2 | 50,2 | 80,0 |
|  | 4 | 14,1 | 32,6 |
| phyDis | 1 | 73,5 | 109,7 |
|  | 2 | 44,6 | 95,6 |
|  | 4 | 11,6 | 52,8 |

**Table S2.**
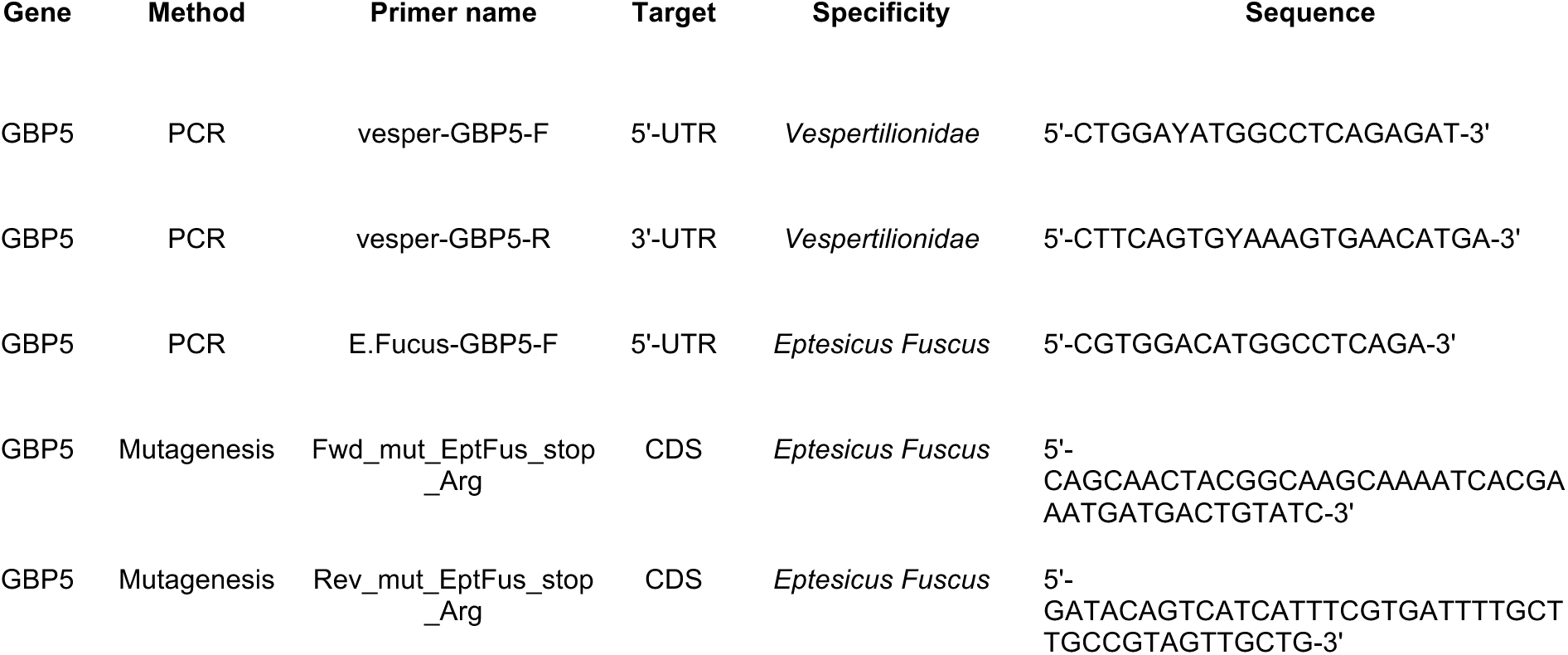
Primer sequences.

